# PANCS-spec-Binders: A system for rapidly discovering isoform– or epitope–specific binders

**DOI:** 10.1101/2025.11.19.689279

**Authors:** Joshua A. Pixley, Matthew J. Styles, Kanokpol Aphicho, Micheal Endres, Priyanka Gade, Karolina Michalska, Andrzej Joachimiak, Bryan C. Dickinson

## Abstract

Proteins that bind to a target protein of interest, termed “binders,” are essential components of biological research reagents and therapeutics. Target proteins present multiple binding surfaces with varying interaction potential. High-potential surfaces, or “hot spots,” are experimentally identified as the most probable binding sites in *de novo* discovery campaigns. However, hot spots and their default binding modes do not always confer the desired specificity. Related proteins or isoforms often share similar hot spots, resulting in promiscuous binding. Interaction with a hot spot may also fail to elicit the intended biological outcome. Consequently, methods that direct *de novo* binder discovery toward targets with defined specificity are critically needed. We recently developed phage-assisted non-continuous selection of binders (PANCS-Binders), a selection platform with unparalleled speed and sequence-function fidelity that enables routine *de novo* binder discovery within days. However, because PANCS-Binder selections enrich variants based primarily on affinity, secondary screening is unlikely to identify binders to lesser hot spots because of the high likelihood of convergence. These alternative binding surfaces with weaker inherent interactions may possess desirable specificity profiles. Here, we develop PANCS-spec-Binders, which incorporates simultaneous selection and counterselection to control the specificity of enriched binders. We demonstrate PANCS-spec-Binders in two proof-of-concept applications: (1) discovery of isoform-selective binders that bind HRAS with >100-fold higher affinity than the highly related KRAS isoform, and (2) discovery of epitope-specific binders that either target or avoid the LIR interaction region of LC3B. PANCS-spec-Binders enables rapid identification of binders with defined specificity within days.

**SIGNIFICANCE:** Affinity reagents, termed “binders”, are essential tools in research and therapeutic development. Binders generally require high specificity either at the selectivity level (binding only the target protein, but not related proteins) or at the epitope level (binding only at a specific surface on the target rather than another). While methods to discover binders in general have progressed, identifying binders with defined specificity features often requires extensive secondary screening and frequently results in failure. Here, we adapt our recently developed binder discovery platform to solve these two selectivity problems. By rapidly screening billions of variants, we can direct specificity between highly related proteins, direct binding to a specific epitope, and, because of the fidelity of our selections, identify binders that specifically avoid a defined epitope.

## INTRODUCTION

Proteins that interact with targets of interest, known as binders, are staples of both basic and translational biological research and have emerged as leading pharmaceutical products in the 21^st^ century(1, 2). These binders can often alter the function, stability, location, interaction partners, or other features of the target protein of interest (POI)(3-6). Binding requires a “hot-spot” on the surface of the target protein: a location with intrinsic capacity for binding due to unfavorable interactions with the solvent (water) and/or the potential for favorable interactions with a binding partner(7-9). Because proteins often have multiple potential “hot-spots,” binders to different surfaces on a target protein can likely be discovered (**Fig. 1A**). However, practical application of binders generally requires that binders make specific interactions, both in terms of selectivity for the target (**Fig. 1B**) and in terms of the epitope of binding to the target (**Fig. 1C**). The degree of this specificity required is determined by the intended application: a hot spot present on two related isoforms is ideal if the desired binder is to interact as a pan-isoform binder, but not if an isoform-selective binder is needed (**Fig. 1B**). Similarly, binding a hot-spot of a native PPI will disrupt the PPI while binding a different hot-spot will preserve the native PPI and allow for simultaneous binding (**Fig. 1C**). This leads to a fundamental tension: binders discovered by methods that are agnostic to binding “hot-spots” will identify binders to the “hottest” spot for a given ligand scaffold, regardless of the desired specificity.

**Fig. 1:**
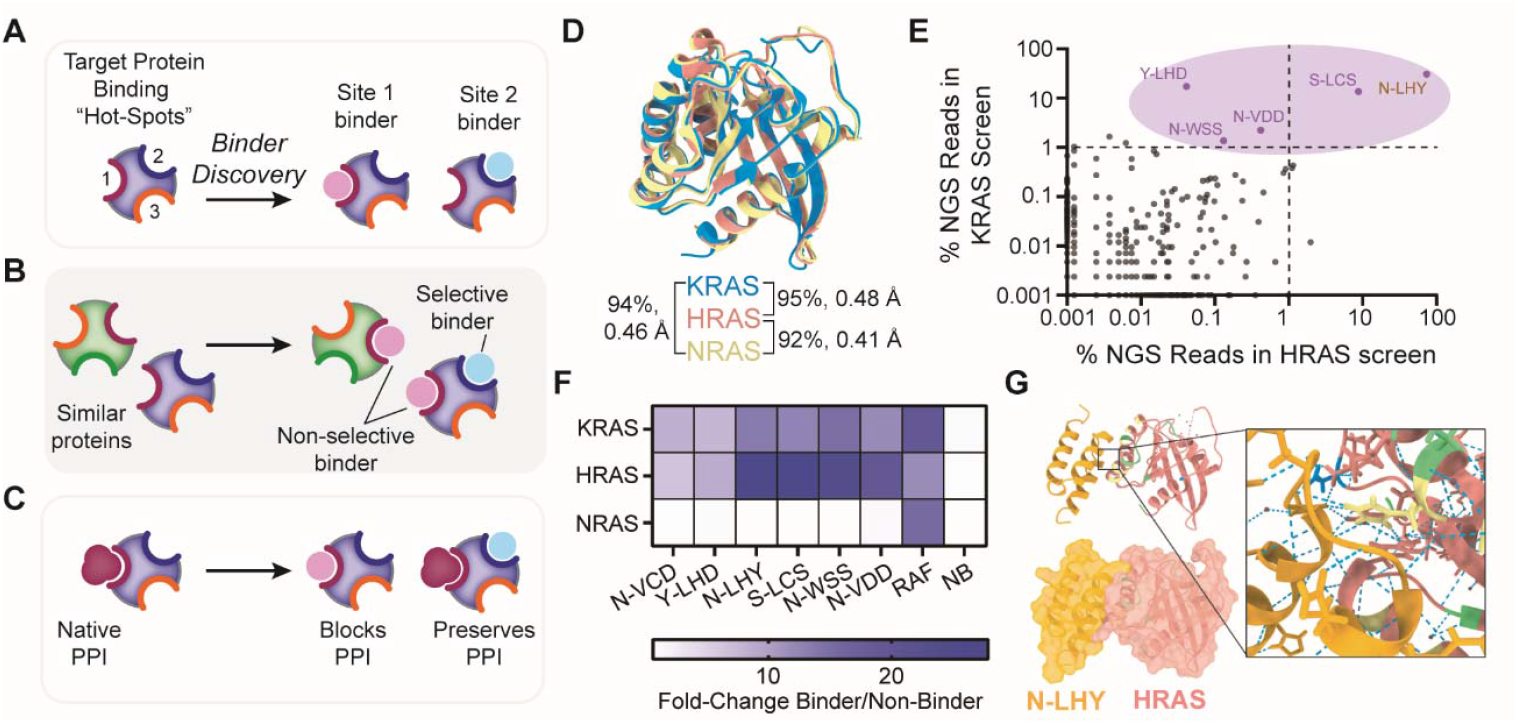
The need for selective binders and Ras as a challenging example. **a**, Binding to a protein occurs at one of several “hot-spots,” and for any given binding scaffold (illustrated here as a circle), binding can occur at multiple sites. **b**, Depiction of binding specificity at the level of isoform selectivity: promiscuous versus selective binders of related proteins. **c**, Depiction of binding specificity at the level of the binding epitope: binding that either disrupts or preserves a native PPI, either of which could be the desired function. **d**, Alignments of crystal structures for the folded regions of KRAS (1-169, blue, PDB: 4obe), HRAS (1-166, red, PDB: 3l8z), and NRAS (1-166, green, PDB: 5uhv) with pairwise sequence (percentage) and structural (RMSD, Å) homology(35-37). **e**, Sequence prevalence for all affibody variants enriched from our 10^10^ library screens for HRAS and KRAS (G12D) - with ZB_neg_ as the counterselection protein on the -AP (33) (n = 6,887). Variants present in only one screen are arbitrarily assigned a prevalence of 0.001% in the opposing screen to facilitate a log-log graph. Correlation analysis shows statistically significant correlation: Pearson r = 0.1178, p < 0.0001 (****). **f**, *E. coli* split RNAP luciferase binding assays for binders isolated from the KRAS PANCS binder selections (**Fig. S1**). **g**, Crystal structure of HRAS (1-166) with N-LHY affibody (PDB ID: 9Z5W), **Table S1**). Residues that differ between HRAS and NRAS are colored yellow on the HRAS structure; likewise, those that differ for KRAS are blue; and those that differ for NRAS and KRAS in green (**Fig. S3** for sequence alignment and **Fig. S4** for more details on the structure.).

Binders are typically discovered through animal immunization to elicit an antibody response(10, 11), various selection platforms such as phage display or FACS (12-14) to mine high-diversity libraries, or more recently, through computational design (15-18). Specific binders, either in the selective or epitope sense, can be discovered through immunization, primarily through extensive screening of antibodies for the desired specificity(19). Display-based selections can be modified to preferentially enrich binders with a desired specificity by alternating selection (for the protein of interest) and counterselection against either a related protein (isoform) or a decoy epitope (20) (mutations on the target protein at the desired binding epitope)(21). Variants that bind in the selection are retained while those that bind in the counterselection are removed. This approach has been applied to a wide variety of specificity problems(5, 21-28). Additionally, masking approaches rely on a known binder, for example, a native PPI partner, to block binding to one epitope in a positive selection, thereby driving enrichment of binding to hot-spots not at that epitope(28). Finally, computational design is privileged in epitope-directed binder discovery because the location of binding can be specified, allowing for the desired specificity to be included in the design process(17, 29-32). However, despite these successes, the discovery of a specific binder, even when successful, generally requires extensive optimization of selection conditions or computational design followed by extensive secondary screening and maturation.

We recently reported Phage Assisted Non-Continuous Selection of Binders (PANCS-Binders)(33), which can screen tens of billions of potential binding variants in just days, dramatically accelerating the binder discovery process. PANCS-Binders uses libraries of replication-deficient phage that encode a potential binding variant fused to half of a proximity-dependent split RNA polymerase (RNAP) and an *E. coli* selection strain that encodes a target protein fused to the other half of the split RNAP and RNAP-controlled expression of an essential phage gene (*gIII*). Phages that encode binders are able to reconstitute the split RNAP to express *gIII* and replicate. In PANCS-Binders, variants are enriched in proportion to their affinity. Highly enriched variants bind and dramatically lower the false positive rate, thereby abrogating the need for secondary screening. Herein, we integrate the principles of differential selections (21) and epitope decoys (20) into PANCS-Binders (34) to establish Phage-Assisted Non-Continuous Selection of Specificity-Oriented Protein Binders (PANCS-spec-Binders), which retains the speed and fidelity of PANCS-Binders but adds a strong counterselection strategy to tune the specificity of enriched binders. We demonstrate the power of PANCS-spec-Binders in two applications: isoform-specific binder discovery using Ras, and epitope-specific binder discovery using LC3B. In both cases, we found that counterselection was able to redirect the selection outcomes of PANCS-Binders to yield protein binders with desired specificity.

## RESULTS

### Identifying RAS Binders from PANCS-Binder Selections

Mutations to the RAS family of proteins are frequent drivers of cancer, making them high-value therapeutic and diagnostic targets(38). Selective binding to RAS isoforms is biophysically challenging because RAS proteins have high structural and sequence homology outside of the disordered C-terminal tail region (94% identical, 98% similar, **Fig. 1D**). Therefore, we sought to use RAS isoforms as a testbed to understand and focus specificity in PANCS-Binder selections. Based on their high structural homology, we predicted that binders selected for either KRAS or HRAS can bind both, and that isoform-selective binders would be challenging to discover. In our initial development of PANCS-Binders, we found that sequence abundance, as measured by next-generation sequencing (NGS), correlated with binder affinity. Therefore, we reasoned we could use the differential enrichment of variants from different RAS isoforms to predict binder selectivity.

As expected, when we performed PANCS-Binders to select for binders to HRAS or KRAS(33), the selections enriched highly similar sets of binders, as shown by comparing the rate of enrichment in the NGS of each selection outcome (**Fig. 1E**). While we don’t know what percent enrichment in any given screen will have >>μM K_d_ affinity (as this depends on the number and relative affinity of the best binders in the selection), we hypothesized that the strong enrichment of Y-LHD (17%) and N-WSS (1.4%) in the KRAS selection and the weak enrichment of these variants in the HRAS screen (0.04% and 0.13%, respectively) suggested that these variants were potentially KRAS selective. To test this hypothesis, we measured binding of the top five variants from the KRAS selection to KRAS, HRAS, and NRAS in our *E. coli* split RNA polymerase (RNAP) complementation luciferase reporter assay (**Fig. 1F, Fig. S1**). All of these variants bound to both HRAS and KRAS, indicating that the hottest spot for affibody binding HRAS and KRAS is indeed highly similar, as is well documented in prior display-based selection results(22, 39). However, none bound NRAS. Within these binders, we observed a clear binding motif (**Fig. S2**) that is predicted to interact with the second (86-102aa) and third (126-137aa) α– helices of KRAS, which is 83% identical and 97% similar with HRAS but only72% identical and 86% similar with NRAS (**Fig. S3**). We determined the X-ray crystal structure of HRAS with one representative binder, “N-LHY” (**Fig. 1G, Table S1, Fig. S4**). The experimental structure strongly correlates with the computationally predicted HRAS:N-LHY structure. We used the structure to identify a point mutation, R97D, that disrupts N-LHY binding to HRAS (**Fig. S5**). These results demonstrate that the dominant hotspots for affibody binding to HRAS and KRAS are highly similar (95% similar or identical for interacting residues, **Fig. S3**), and that we were unable to simply use the relative enrichment rate of variants in the two different screens in order to identify highly selective variants, as even Y-LHD, which enriched to only 0.04% in the HRAS selection, still displayed strong binding to HRAS. We suspect that each selection has a different cut-off above which measurable binding occurs (0.1%, 0.01%, etc.). We hypothesize that lowly enriched variants in the non-selective screen might be isoform-specific (by interacting with a lesser hotspot), and therefore, we sought to redesign the PANCS-binder selection system to incorporate a strong counter-selection to assess whether variants with novel specificities could be identified by re-screening the library in a manner that favored HRAS-selective binding.

### Development of PANCS-spec-Binders

We envisioned a modified version of PANCS-Binders method in which we use active counterselection to de-enrich promiscuous variants in isoform selections (**Fig. 2A**). PANCS-Binders uses replication-deficient phage that encode protein libraries tagged with one half of a proximity-dependent split-RNA polymerase (RNAP_N_) biosensor and *E. coli* selection strains engineered to express a target protein of interest tagged with the other half of the split RNA polymerase (RNAP_C-CGG_). Protein-protein interaction (PPI) between a phage-encoded variant and the target reconstitutes the RNA polymerase (RNAP_CGG_) and triggers expression of a required phage gene, *gIII*, allowing the phages encoding that variant to replicate. To prevent variants that cheat the system by binding to the RNAP_C-CGG_ rather than to the target protein from enriching, we utilize an orthogonal RNAP_C-T7,_ which selectively drives expression of a dominant negative variant of *gIII* (*gIII*_*neg*_) that poisons phage growth. In our original work, RNAP_C-T7_ was fused to a zipper peptide, ZB_neg_, used in the evolution of the proximity-dependent biosensor using the split RNAP components(40). We have previously utilized counterselection in continuous evolution (PACE) experiments with this same biosensor to direct the evolution of specificity to convert a promiscuous binder into a selective binder(41). In this strategy, a counterselection protein is displayed on an RNAP_C-T7_ such that promiscuous binding poisons phage replication (**Fig. 2A**). We sought to utilize this capacity for active counterselection during *de novo* selections to drive the specificity of enriched binders, thereby establishing our new platform, PANCS-spec-Binders. To construct our biosensor (**Fig. 2A**), we used our HRAS +AP and replaced the zipper peptide on our –AP with KRAS (fused to RNAP_C-T7_). We then validated that this –AP prevents activity-dependent replication (**Fig. S6**) of phage encoding RNAP_N_-binders that were promiscuous (binding both HRAS and KRAS).

**Fig. 2:**
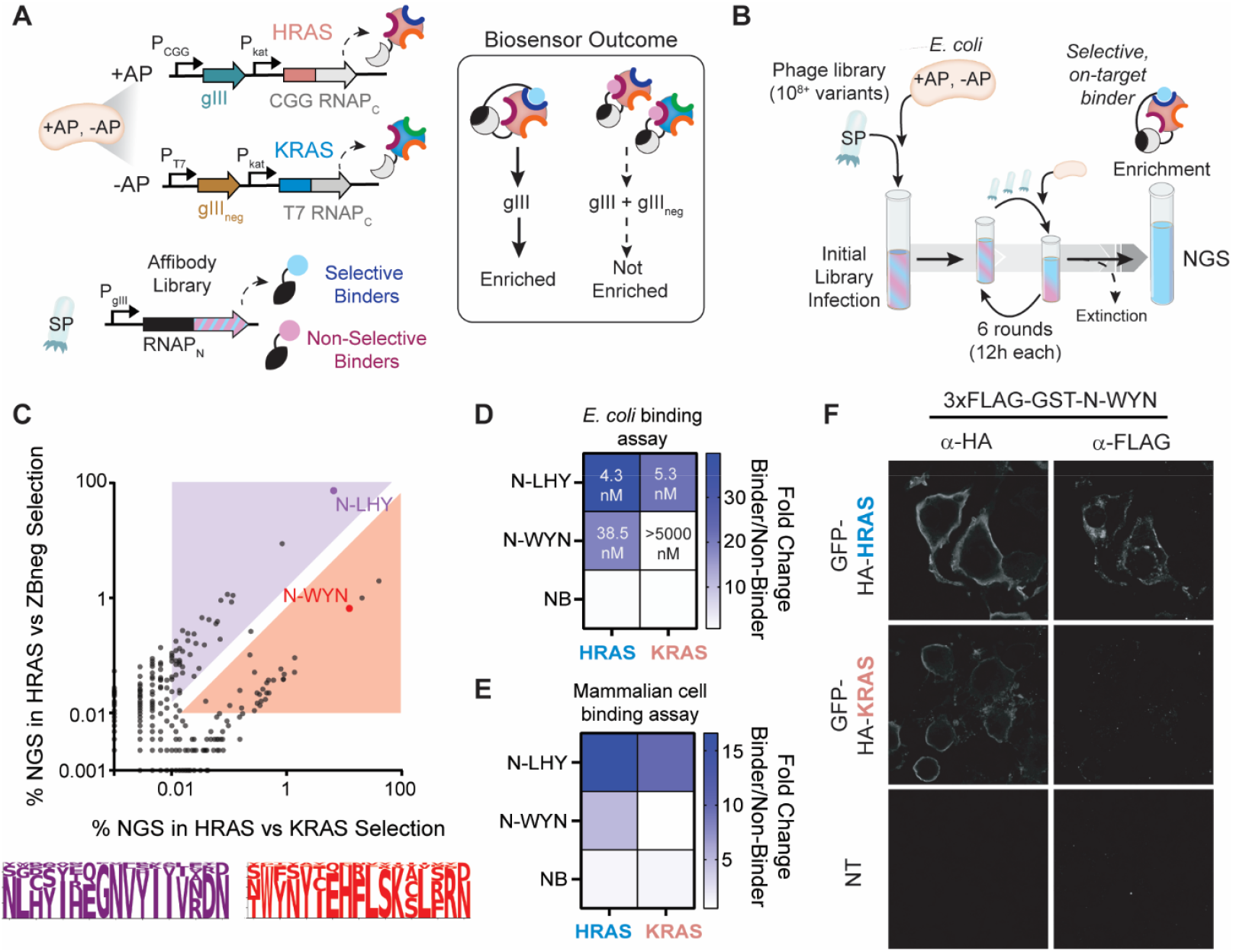
Discovery of an Isoform Selective HRAS Binder. **a**, PANCS-spec-Binders modified biosensor for HRAS isoform-specific binder selection. **b**, Depiction of the PANCS-spec-Binder workflow. **c**, Comparison of the relative enrichment of variants in the selective screen (x-axis), HRAS vs. KRAS, and the non-selective screen (y-axis), HRAS vs. ZB_neg_. If a variant was not present in the NGS of one selection, it was listed as 0.001%. The purple shading and sequence logo represent putative non-selective HRAS binders, and the red shading and sequence logo represent putative selective HRAS binders (**Fig. S8**). **d**, *E. coli* split RNAP luciferase assay (**Fig. S1**) for select isolated binders (**Fig. S10**; circled in **c**). Numerical values indicate K_d_ (measured by SPR, **Fig. S11, Table S3**). **e**, Split-luciferase binding assay in HEK293T cells via transfection of RAS isoform fused c-terminally to the C-terminal portion of split-Nano luciferase and binders fused N-terminally to the N-terminal portion of split Nano luciferase (**Fig. S10**). **f**, HEK293T cells were transfected with GFP-HA-RAS isoforms, and purified 3xFLAG-GST-N-WYN was used as the primary antibody for immunofluorescence imaging (**Fig.S12**). NT: non-transfected controls.

### PANCS-spec-Binders Proof-of-Concept: Discovery of Isoform-Selective RAS Binders

Using a 10^10^ affibody library(33), we ran four parallel selections, each with 6 rounds of serial passaging (**Fig. 2B**): 1) HRAS vs. ZB_neg_, 2) HRAS vs. KRAS, 3) NRAS vs. ZB_neg_, and 4) NRAS vs. KRAS. The endpoint titers of each selection were all above 10^7^ plaque-forming units (PFU)/mL (**Table S2, Fig. S7**), suggesting successful enrichment of binders(33). As a negative control, we also ran a selection of KRAS vs. KRAS, which went extinct, further confirming the PANCS-spec-binder platform can enrich binders with defined specificity.

Comparing the binders enriched in the HRAS vs. ZB_neg_ (all HRAS binders) and HRAS vs. KRAS (isoform-selective HRAS binders) screens, we observed a distinct bifurcation in the enriched variant populations (**Fig. 2C**, purple for promiscuous and red for selective), each with a distinct amino acid profile (**Fig. 2C, Fig. S8**). Conversely, the NRAS selections against ZB_neg_ and against KRAS produced nearly identical enriched variant populations (**Fig. S9)** suggesting that for the affibody scaffold, the hottest spot on NRAS is a surface not present on KRAS (consistent with the lack of KRAS binders with affinity to NRAS, **Fig. 1F**). We isolated a variant (N-WYN) from our HRAS vs. KRAS selective screen and confirmed that N-WYN binds HRAS with no detectable KRAS binding (**Fig. 2D**). To confirm this finding, we determined the affinity of N-LHY and N-WYN for KRAS and HRAS via surface plasmon resonance (SPR). The non-selective binder, N-LHY, displayed high affinity to both (~5 nM K_d_), while the selective binder, N-WYN, showed high affinity to HRAS (39 nM) but no binding to KRAS (tested up to 5 μM; >100-fold selectivity; **Fig. 2D, Fig. S11, Table S3**). These results were confirmed via both split nano-luciferase complementation assay in human HEK293T cells (**Fig. 2E, Fig. S10**) and by immunofluorescence staining of HEK293T cells expressing either HRAS or KRAS (**Fig. 2F, Fig. S12**). Collectively, these selections identified NRAS-specific binders, HRAS-specific binders, HRAS/KRAS-specific binders, and HRAS/NRAS-specific binders (**Table S4, Fig. S13**) and demonstrate that counterselection can be used in PANCS-spec-Binders to preferentially enrich isoform-selective binders. Next, we sought to understand the molecular basis for the rare, HRAS-selective binding mode.

### Elucidating the mode of binding of HRAS selective binders

Despite decades of work discovering binders to the RAS family of proteins, isoform-selective binders have been incredibly challenging to discover(29). Given that PANCS-spec-Binders was able to solve this longstanding biophysical puzzle in a matter of days, we next sought to elucidate the binding mode that facilitated the selective HRAS interaction. First, we predicted the interaction complex for HRAS and our selective binder using AlphaFold3, and identified mutations aimed at disrupting binding (**Fig. 3A**). None of the three mutations tested for disruption of the HRAS:N-WYN interaction resulted in any perturbation to binding (**Fig. 3B, S14**). We also note that the R79D mutation capable of ablating N-LHY binding does not impact N-WYN binding, providing further evidence of an alternative binding site (**Fig. 3B, S14**). We concluded the AlphaFold3 binding prediction was incorrect and sought to experimentally map which amino acids that differ between HRAS and KRAS confer loss of binding to N-WYN. We swapped individual mutations in the structured region of HRAS (Q95H, D107E, A121P, A122S, E126D, S127T, R128K, Y141F), but none of these perturbed N-WYN binding (**Fig. S15**). We then swapped groups of these mutations (**Fig. 3C**), including the entire set of residues in the structured region (95-153); however, these also failed to perturb the HRAS:N-WYN interaction (**Fig. 3D, Fig. S16**). Finally, we investigated whether the disordered C-terminal tail of HRAS was involved in binding. Swapping the C-terminus (165-188) from KRAS onto HRAS conferred loss of binding, suggesting that the disordered tail may be the site of binding (**Fig. 3D**).

**Fig. 3:**
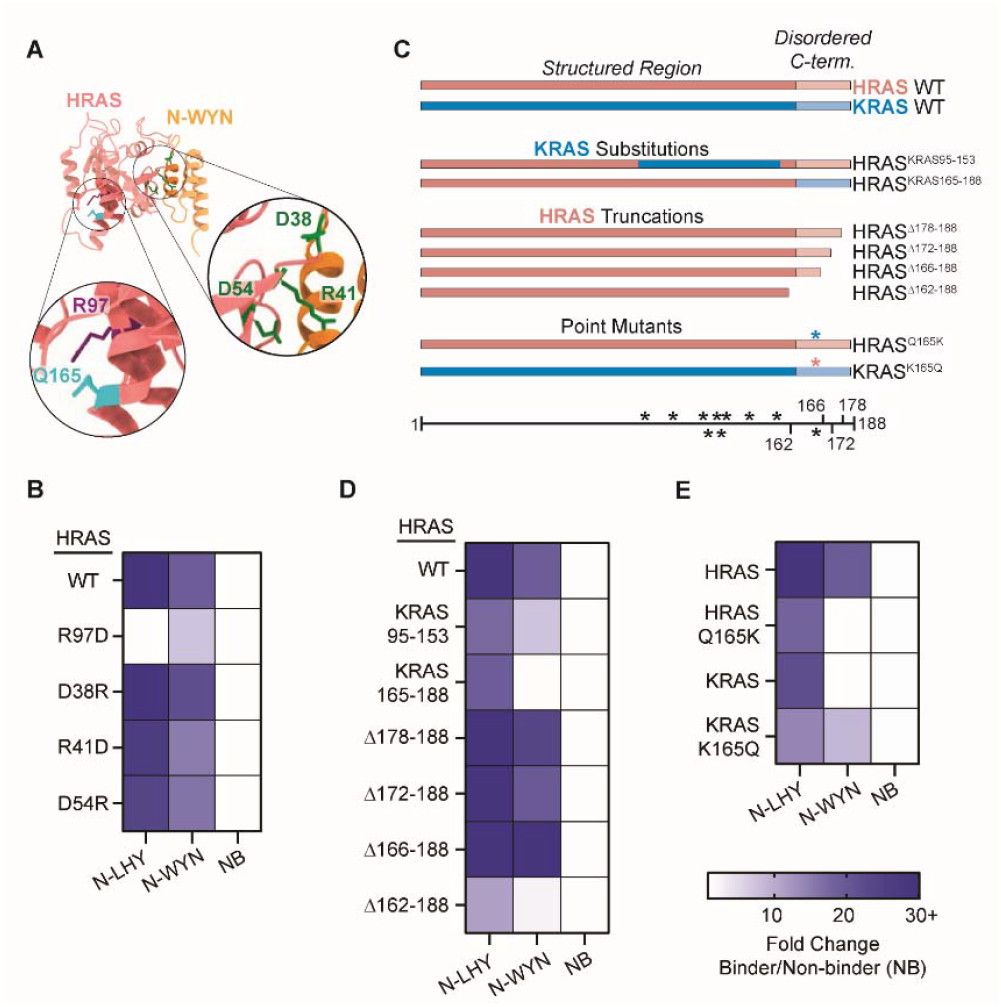
Mechanism of HRAS Selective Binding. **a**, AlphaFold3-predicted N-WYN (gold):HRAS (red) complex. Selected positions for mutations on HRAS are shown as colored spheres: R97 (purple, disrupts N-LHY binding); D38, R41, and D54 (green, predicted to disrupt N-WYN binding); and Q165 (cyan, disrupts N-WYN binding). **b**, *E. coli* luciferase assays for N-LHY and N-WYN to HRAS mutants selected based on the AlphaFold3 predicted structures for N-LHY and N-WYN (**Fig. S14**). **c**, A cartoon depicting the position of HRAS to KRAS mutations and HRAS truncations tested in our *E. coli* luciferase assay (**Fig. S16)**: * indicate positions where HRAS and KRAS sequence differs, red indicates the HRAS sequence was used, and blue indicates the KRAS sequence was used. The lighter color indicates the disordered C-terminal tail. **d**, *E. coli* luciferase assays for N-LHY and N-WYN to HRAS mutant and truncations highlighted in **c. e**, Same as **d** with Q165 (cyan in **a**) identified as a mutation that confers N-WYN binding (**Fig. S16**). The legend shown below **e** applies to **b, d**, and **e**. In each case, NB indicates a non-binding affibody.

We next tested several truncations on HRAS, showing that Δ167-188 (loss of the entire tail region commonly excluded in HRAS purification (HRAS Δ170-188), including in our SPR experiments) retained N-WYN binding, but Δ163-188 lost binding. HRAS and KRAS differ at only one position between 163-166: K165 for KRAS and Q165 for HRAS **(Fig. S3)**. A single point mutation at this position converts N-WYN into a KRAS binder and disrupts binding to HRAS (**Fig. 3E, Fig. S16**). Interestingly, NRAS has Q165 like HRAS, but is different at the adjacent residue (HRAS is QH, NRAS is QY, and KRAS is KH at 165/166), but N-WYN did not enrich well in the NRAS vs zipper peptide selection (0.01%) (**Fig. S17)**. Interestingly, N-WYN shows some binding to NRAS in our *E coli* binding luciferase assay, but the related S-WFS from the HRAS vs KRAS selection does not (**Fig. S13**). These results suggest the presence of an HRAS-specific hot spot at this position.

### PANCS-spec-Binder selections to direct the epitope of binding

Having demonstrated the capability of PANCS-spec-Binders to enrich isoform-specific binders, we next sought to direct specificity to different surfaces within a single target (epitope specificity, **Fig. 1C**). In our previously reported selections, we assessed pairwise binding specificity of 15 *de novo* binders, which were all very selective, except for the LC3B binder, which also bound to GABARAP. While having low overall sequence similarity (**Fig. 4A**), GABARAP and LC3B are members of a broader family (42) of proteins that interact with LIR motifs to recruit proteins into the autophagosome(43). Indeed, our LC3B binder appears to have an LIR motif (FEIL) enabling it to bind to both LC3B and GABARAP (**Fig. 4B**). Notably, the GABARAP-selective binder does not contain this motif, likely pointing to the presence of a separate, GABARAP-specific hot-spot. LC3B has been the focus of several targeted-protein degradation strategies, including AUTACSs, ATTECs, ATNCs, etc., wherein molecules induce proximity between LC3B and a target protein to facilitate target degradation via autophagy(44-46). Most of these techniques have relied on LIR motifs or mimics for binding to LC3B(47). As these interactions compete with native LIR interactions and interact with other LIR-interacting proteins(43), we sought to identify a new binding hotspot on LC3B outside of the LIR-binding site.

**Fig. 4:**
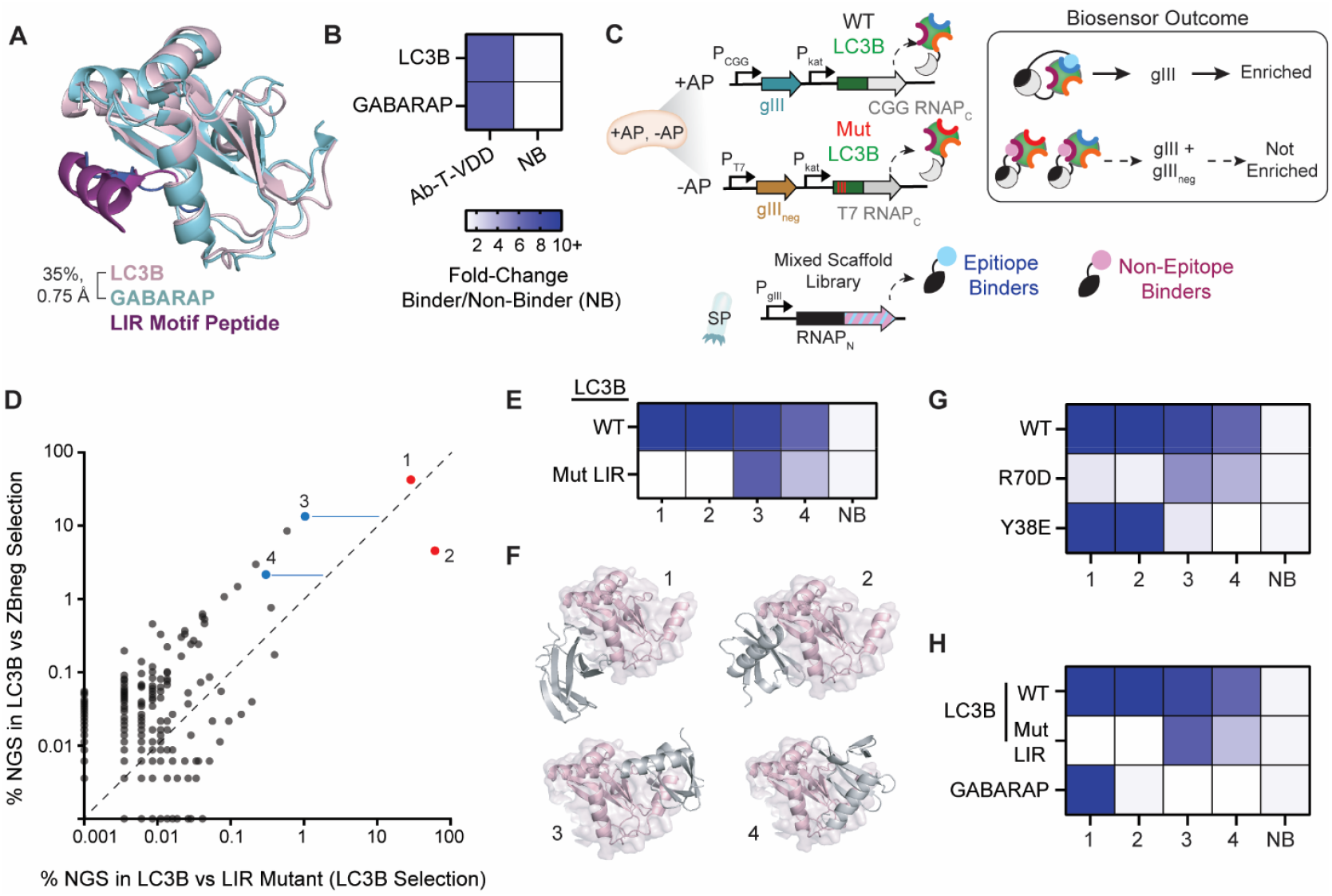
Directing Epitope Specificity of Binder Discovery. **a**, Structure of LC3B (pink, PDB: 5WRD) and GABARAP (teal, PDB: 4XC2) in complex with respective LIR-motif containing peptides (purple and blue). **b**, *E. coli* lux binding data for previously published promiscuous binder to LC3B and GABARAP. **c**, Workflow for the generation of epitope-selective protein binders using PANCS-spec-Binders selections. **d**, Comparison of the relative enrichment of variants in the selective screen (x-axis), LC3B vs. LC3B LIR Mutant, and the non-selective screen (y-axis), LC3B vs. zipper peptide. If a variant was not present in the NGS of one selection, it was listed as 0.001%. **e**, *E. coli* split RNAP luciferase assay for select isolated binders (circled in **d**; **Fig. S21**). **f**, Alphafold3-predicted complexes of each isolated binder with LC3B show potential preference for two distinct binding faces. **g**, *E. coli* lux binding data for our isolated binders with mutations selected to disrupt binding at one of the two hot-spot surfaces (**Fig. S22**). **h**, *E. coli* lux of each isolated binder with LC3B, LC3B with mutations at the LIR, and GABARAP (**Fig. S24**).

To enforce epitope-specificity, we used an epitope-decoy strategy with PANCS-spec binders: the counterselection protein was engineered to be identical to the target except for mutations within the desired binding epitope. This makes non-epitope binding hot spots the same on both proteins, enriching binders that recognize only the intended epitope (**Fig. 4C**). Zhou *et al*. recently developed a set of heuristics for identifying residues to generate epitope decoys for counterselections in phage display, including targeting large polar residues and residues on loops(20). As we did not know the location of the second “hot-spot” *a priori*, we sought to identify binders to an alternative site indirectly, by inference. We aimed to identify all LC3B binders in a non-directed selection and all LIR binders in an LIR site-directed selection; those relatively de-enriched in the epitope-directed selection are likely to bind to a different epitope. To direct binding only to the LIR region of LC3B, we placed LC3B WT on the +AP and a mutant version of LC3B on the –AP, such that only binders that interact with LC3B WT at the same place as the mutation sites on the -AP would replicate (**Fig. 4C**). We identified several surface exposed residues of LC3B that interact with an LIR peptide (PDB 5wrd) and mutated them to present a different surface on the -AP: E62K, K65D, I66S, R69E. This set of mutations produced a decoy that expressed similarly well as the wild type (**Fig. S18**).

To maximize our chances of identifying binders to an alternative hot spot, we used a mixed scaffold library for our selections, including affibodies, affitins, nanobodies, and monobodies (**Fig. S19**)(27, 48-50). As a negative control, we also ran a selection with LC3B WT on the +AP and -APs, which went extinct after 4 passages, while our selection conditions retained high titers (**Table S5**, NGS in **Fig. S20**). When comparing the results of our promiscuous and selective selections, one population of binders was heavily enriched in both selections (LC3B WT vs ZB_neg_ and LC3B WT vs LC3B Mutant LIR; **Fig. 4D**, along the diagonal), while a second population of binders was de-enriched in the epitope-directed conditions (LC3B WT vs LC3B Mutant LIR) compared to the non-directed conditions (LC3B WT vs ZB_neg_; **Fig. 4D**, left-shifted from diagonal). Based on these enrichment patterns, we selected two binders from the first category and two from the second (these latter binders had 12.9- and 7.8-fold de-enrichment in the selective screen relative to the non-selective screen, blue lines in **Fig. 4D**) and confirmed the expected binding pattern (**Fig. 4E, Fig. S21)**. The AlphaFold3-predicted interaction complex for LC3B with these binders suggested two independent binding surfaces for the two binder populations (**Fig. 4F**). Based on the predicted structures, we identified a set of single-point mutations that selectively disrupt binding at these two surfaces, thereby identifying binders that do not interact with the LIR binding region of LC3B, as desired (**Fig. 4G, Fig. S22**). Interestingly, further investigation showed that the determined binding site is close to another protein-protein interaction surface between LC3B and ATG4b protease, which cleaves LC3B to create a new C-terminus prior to lipidation (**Fig. S23**)(51). Finally, since our original LC3B binder discovered through PANCS-Binders was promiscuous with GABARAP, we checked whether our new set of binders was specific for LC3B. As expected, given the low sequence similarity outside of the LIR binding region, binders that interact with the new site are specific, and one of the two LIR-like binders is specific for LC3B (**Fig. 4H, Fig. S24**). This is not unexpected, as several LIR motifs exist for which there is some specificity among these homologs(52, 53). In all, we demonstrate that PANCS-spec-Binders selections can be used to direct the epitope specificity of enriched binders, and because of the high fidelity of the selections, we can inferentially identify new binding sites when the desired epitope of binding is unknown/unspecified.

## DISCUSSION

In binder discovery, the intrinsic compatibility of a surface of a protein for binding interactions governs whether or not identification of a binder (in the context of a given library) can be performed. Often, the surfaces compatible with binding (hot spots) are not known *a priori*, but rather, are determined experimentally. For many applications, especially therapeutic ones, binding must be to a specific epitope (for example, to disrupt a PPI, **Fig. 1C**). Therefore, methods to direct binding among the available hot-spots towards specific hot-spots are often necessary. PANCS-spec-Binders offers several key advantages that allow for rapid selective binder discovery: 1) multiplexing capability to test many counterselection decoy epitopes at once, 2) facile tunability of the counterselection pressure via multiplexing varied expression levels of the same decoy epitope to enrich binders with moderate selectivity, and 3) output binders can be used directly without extensive screening due to the extremely high fidelity of PANCS selections. This fidelity even enables identification of binders to unknown hot-spots by inference between the population of all binders to a target protein and of binders specifically to a known binding epitope of a target protein.

Binder discovery is increasingly becoming a solvable problem with the improvement of computational and experimental methods. However, this work demonstrates that minor modifications to a single side chain (Q vs K) can be critical for determining selectivity; additionally, while AlphaFold3 was able to predict RAS isoform specificity for one binding motif (the HRAS/KRAS motif in **Supplementary Fig. 2**), it was unable to predict the specificity for either the HRAS or the NRAS specific binders suggesting that AlphaFold3’s performance is highly bifurcated – predicting certain hotspot interactions very well while completely failing at others (**Fig. S25**). While we focused in this study on binder specificity at the isoform- and epitope-level, PANCS-spec-Binders is suitable for directing the specificity of binders in all cases where the epitope of interest and decoy epitope of interest can be genetically encoded in the context of the *E. coli* cytoplasm. This includes mutant, splicing variant, and proteolytic cleavage-specific binders, as these can be directly genetically encoded as we have performed here.

Additionally, using non-canonical amino acid incorporation via amber suppression, various post-translational modifications could be incorporated either into the epitope of interest or the epitope decoy (phosphoserine(54), phosphotyrosine(55), methyllysine(56), etc.(57)). Our test cases, isoform and epitope selectivity, demonstrate the speed and utility of PANCS-spec-Binders for directing the specificity in binder discovery campaigns.

## Materials and Methods

Full details regarding materials and methods are provided in the Supplemental Materials. Generally, cloning, plaque assays, selections, luciferase assays, SPR, and NGS analysis were performed as previously described(33). Select changes are described below.

### PANCS-spec-Binder Selections

Briefly, a selection strain (+AP/-AP) is grown to the stationary phase in LB with carbenicillin and kanamycin, subcultured 1/10 in fresh LB with carbenicillin and kanamycin to an OD600 of 0.4-0.6 prior to adding phage. For passage 1, stock phages are added to the subculture and incubated at 37 °C with shaking (200 rpm) for 12 h, then centrifuged to pellet the cells and collect the cell-free supernatant (referred to as passage 1 phage). For subsequent passages, a fraction (5%) of the cell-free supernatant from the prior passage is added to the subculture and incubated at 37 °C with shaking (200 rpm) for 12 h, then centrifuged to pellet the cells and collect the cell-free supernatant (referred to as passage # phage). For the 10^10^ affibody library PANCS, 6 passages were performed with a 5% transfer rate between passages. Passage 1 was seeded with 5*10^10^ PFU in a 500 mL subculture.

Smaller subcultures were used for subsequent passages: 125 mL for passage 2, 25 mL for passage 3, and 5 mL for passages 4-6. For the mixed scaffold library, 4 passages were performed with a 5% transfer rate between passages. Passage 1 was seeded with 2*10^9^ PFU of the pooled library (**Fig. S19**) in 5 mL of subculture. The transfer rate (5%) and subculture volume (5 mL) were both held constant throughout the selection. In both cases, titers of the final passage were then determined using activity-independent plaque assays.

### Co-expression and Purification of HRAS (1–166)/N-LHY Affibody Complex

The genes encoding HRAS (1–166) and N-LHY affibody were cloned into the expression vectors pMCSG53 and pMCSG120, respectively, which share identical multiple cloning sites. The vector pMCSG53 carries ampicillin resistance and a ColE1 origin of replication, whereas pMCSG120 confers kanamycin resistance and contains an RSF origin of replication. The HRAS (1–166) construct included an N-terminal His_6,_ tag followed by a tobacco etch virus (TEV) protease cleavage site. For co-expression, HRAS (1–166) _pMCSG53 and N-LHY affibody_pMCSG120 plasmids were co-transformed into *Escherichia coli* BL21(DE3)-Gold cells (Stratagene). Cultures were grown in LB medium supplemented with ampicillin (150 μg/ml) and kanamycin (100 μg/ml) at 37 °C until the optical density at 600 nm reached 1.0. Protein expression was induced with 0.4 mM isopropyl β-D-1-thiogalactopyranoside (IPTG), followed by incubation at 18 °C for 16 h. Cells were harvested by centrifugation and resuspended in buffer A (50 mM HEPES/NaOH, pH 8.0, 0.5 M NaCl, 20 mM imidazole, 5% glycerol, and 10 mM β-mercaptoethanol). Cell disruption was achieved by sonication (5 min total, 130 W output), and the lysate was clarified by centrifugation at 30,000 × g for 1 h at 4 °C. The Ni^2+^ affinity purification step was performed using a Flex-Column connected to a Van-Man vacuum manifold (Promega). Briefly, supernatant was loaded on 5 ml Ni^2+^ Sepharose (GE Healthcare Life Sciences) equilibrated with buffer A and mixed thoroughly with the resin. A vacuum of 15 psi was used to speed the removal of supernatant and to wash out unbound proteins (150 ml buffer A supplemented to 50 mM imidazole). The bound proteins were eluted with buffer A supplemented with 500 mM imidazole. The eluted complex was dialyzed against buffer A, and the His_6,_ tag of HRAS (1–166) was removed by incubation with TEV protease at a 1:20 protease-to-protein molar ratio for 16 h at 4 °C. The cleavage mixture was passed through a Ni-NTA gravity column to remove His-tagged TEV protease and residual contaminants. The flowthrough containing the HRAS (1–166)/N-LHY affibody complex was concentrated and further purified by size-exclusion chromatography on a Superdex 75 10/300 GL column (GE Healthcare) equilibrated with 20 mM HEPES/NaOH, pH 7.5, 150 mM NaCl, and 1 mM TCEP. Fractions containing the complex were pooled, concentrated to 35 mg/ml, and flash-cooled in liquid nitrogen and stored at –80 °C.

### Crystallization, X-ray diffraction data collection and structure solution

The HRAS (residues 1– 166)/N-LHY affibody complex was crystallized using the sitting-drop vapor diffusion method at 16 °C. Each crystallization drop contained 0.4 μl of the protein complex solution mixed with 0.4 μL of reservoir solution (0.2 M ammonium nitrate, pH 6.3, and 20% (w/v) PEG 3350). Crystals of the HRAS (1–166)/N-LHY affibody complex were cryoprotected in reservoir solution supplemented with 20% (v/v) ethylene glycol and subsequently flash-cooled in liquid nitrogen. X-ray diffraction data were collected at beamline 19-ID of the National Synchrotron Light Source II (NSLS-II) at Brookhaven National Laboratory. The diffraction images were recorded on the Eiger2 9M XE detector and processed using the Xia2 automated data-processing pipeline(58). The structure was solved by molecular replacement with the MolRep program (59) using an AlphaFold3 model (60) of HRAS (1–166)/N-LHY affibody complex as a search model. The initial model was refined through iterative model building in COOT (61) and crystallographic refinement in Phenix (62) and Refmac(63).

### Immunofluorescence staining using purified recombinant NWYN binder and confocal microscopy

#### Plating and transfection

HEK293T cells were plated on glass coverslips (Neuvitro Corporation; GG-12-15-oz) pre-treated with 0.05 mg mL-1 poly-D-lysine (Sigma-Aldrich; P7280) in a 24-well dish. At 18-24 h post-plating, The cells were transfected with 500 ng plasmid or an equal volume of water using Lipofectamine 3000 Transfection Reagent (Invitrogen; L3000008) in 500 uL Opto-MEM reduced serum medium (Gibco; 31985070), according to manufacturer’s instruction. The medium were replaced with complete medium (DMEM containing 10%FBS) at 3-4 h post transfection.

#### Fixation, Permeabilization, and Staining

At 18-24 h post transfection, cells were fixed with 4%paraformaldehyde in PBS (freshly prepared from 16%paraformaldehyde; Electron Microscopy Sciences; 15710) at room temperature for 15 minutes and then permeabilized with cold methanol at -20 ° C for 7 minutes. The samples were blocked with SuperBlock Blocking Buffer in PBS (Thermo Scientific; 37515) at room temperature for 30 minutes and then incubated with or without 3xFLAG-GST-NWYN (500 nM) at room temperature overnight. The samples were simultaneously incubated with the primary antibodies at room temperature for 1 h: Rabbit Anti-HA (1:1000 dilution, Cell Signaling Technology; 3724) and Mouse Anti-FLAG (1:1000 dilution, Sigma-Aldrich; F3165). The samples were then simultaneously incubated with the secondary antibodies at room temperature for 1 h: Anti-Rabbit-AF568 (1:1000 dilution, Invitrogen; A-11011) and Anti-Rabbit-AF647 (1:1000 dilution, Invitrogen; A32728). Finally, nuclear counterstaining was performed using 1 μM DAPI in PBS for 3 min. Unless otherwise stated, cells were washed twice with PBS between steps, and SuperBlock in PBS was used for antibody and protein dilutions.

#### Mounting

The coverslips were transferred out of the 24-well plate and inverted onto a drop of a mountant (ProLong diamond antifade mountant, ThermoFisher; P36961) on a glass slide (Fisherbrand; 12-544-2). The slides were allowed to cure for at least 24 hour in the dark before imaging.

#### Fluorescence confocal microscopy

Confocal imaging was performed on a Leica SP8 laser-scanning confocal microscope at the University of Chicago Integrated Light Microscopy Facility (RRID: SCR_019197). All images were acquired using an HC PL APO CS2 63×/1.40 NA oil-immersion objective with or without a 2.5× zoom factor. Images were collected in 16-bit format at a 2048 × 2048-pixel resolution, with the pinhole set to 0.75 Airy units, and acquired at a scan speed of 400 Hz in line-sequential mode with 4× line averaging. The following sequential configuration was used:

DAPI channel: 405 nm diode laser at 1 % intensity; detection via PMT (410–490 nm), gain 500.

EGFP channel: 488 nm line from the white-light laser at 0.2 % of 70 % maximum power; detection via HyD SMD (493–572 nm), gain 20.

AF568 channel: 577 nm white-light laser at 1 % of 70 % maximum power; detection via HyD SMD (582– 648 nm), gain 100.

AF647 channel: 653 nm white-light laser at 1 % of 70 % maximum power; detection via HyD (658–776 nm), gain 100.

#### Fluorescence image quantification and analysis

Acquired images were visualized and analyzed using Fiji (PMID: 22743772 and PMID: 22930834). Background and contrast were adjusted uniformly across all images. Background adjusted images were exported as 8-bit grayscale for quantification of mean fluorescence intensity. All images are shown in **Fig. S26** and **S27**.

## Acknowledgements

This work was supported by the National Institute of General Medical Sciences (GM119840 to B.C.D and F32GM147968 to M.J.S) and then National Cancer Institute (P30CA014599) of the National Institutes of Health, and by the Camille and Henry Dreyfus Foundation Teacher Scholar Award (B.C.D.). The use of SBC resources at the Advanced Photon Source is supported by the U.S. Department of Energy (DOE) Office of Science and operated for the DOE Office of Science by Argonne National Laboratory under Contract No. DE-AC02-06CH11357. We thank S. Ahmadiantehrani for assistance with preparing this paper.

## Contributions

Conceptualization: J.A.P., M.J.S. and B.C.D. Methodology: J.A.P. Investigation: J.A.P.; M.J.S.; and K.A.. Crystallography: M.E., P.G., K.M, and A.J. Writing: J.A.P., M.J.S., and B.C.D. Supervision: B.C.D.

## Ethics Declaration

Competing interests B.C.D. is an inventor on the patent describing the split RNAP biosensors. The University of Chicago has filed a provisional patent on the PANCS-Binders technology with M.J.S and B.C.D. listed as inventors.

## SUPPORTING INFORMATION

Details of bacterial strains, plasmids, primers, additional data figures and tables.

