## Supporting Information for "PANCS-spec-Binders: A system for rapidly discovering isoform– or epitope–specific binders"

**TABLE OF CONTENTS**

**Supplementary Fig. 1: *E. coli* Binder Lux Assay of Binders Isolated from PANCS-Binder Selections**

**Supplementary Fig. 2: Affibody Motif for Binding KRAS**

**Supplementary Fig. 3: Sequence Alignment of RAS Isoforms Aligned to AF3 Binding Mode**

**Supplementary Table 1: NLHY-HRAS Crystal Structure Details**

**Supplementary Fig. 4: Additional Details of N-LHY-HRAS Binding Interaction**

**Supplementary Fig. 5: Mutation that disrupts N-LHY binding to HRAS**

**Supplementary Fig. 6: Optimization of KRAS -AP Expression Level**

**Supplementary Table 2: RAS PANCS-spec-Binder Selection Endpoint Titers**

**Supplementary Fig. 7: RAS Selection Endpoint NGS**

**Supplementary Fig. 8: Sequence Logos of Selection Endpoints and Lux to Confirm Binding**

**Supplementary Fig. 9: NRAS Selection Enrichment Comparison and Lux Validation**

**Supplementary Fig. 10: Individual Lux data for 2E and 2F**

**Supplementary Fig. 11: SPR Plots for Selected Binders**

**Supplementary Table 3: Kinetic Fits to SPR Plots for Selected Binders**

**Supplementary Fig. 12: IF Replicate Images and Quantification**

**Supplementary Table 4: Summary of RAS Specific Binder Variants**

**Supplementary Fig. 13: Cross Reactivity of Isoform Specific RAS Binding Variants**

**Supplementary Fig. 14: Individual Lux data for 3B**

**Supplementary Fig. 15: Lux of Individual Point Mutations to Swap KRAS Mutations into HRAS**

**Supplementary Fig. 16: Individual Lux data for 3C and 3D**

**Supplementary Fig. 17: Comparison of HRAS vs $ZB_{neg}$ and NRAS vs $ZB_{neg}$**

**Supplementary Fig. 18: Expression Level of LC3B and LC3B LIR Mutant**

**Supplementary Fig. 19: Mixed Scaffold Library Details**

**Supplementary Table 5: LC3B Selection Endpoint Titers**

**Supplementary Fig. 20: NGS of LC3B Selections**

**Supplementary Fig. 21: Individual Lux data for 4E**

**Supplementary Fig. 22: Individual Lux data for 4G**

**Supplementary Fig. 23: LC3B:ATG4b Interaction Site**

**Supplementary Fig. 24: Individual Lux data for 4H**

**Supplementary Fig. 25: AlphaFold3 Fails to Predict Binder Specificity**

**Supplementary Fig. 26: Confocal microscopy demonstrated HRas-selective immunofluorescence staining using purified recombinant N-WYN binder**

**Supplementary Fig. 27: Zoom-out micrographs corresponding to 1× confocal zooms of the fields of view (FOVs) shown in Figure S26**

**Additional Materials and Methods**

**Supplementary Table 6: Plasmids**

**Supplementary Table 7: Primers**

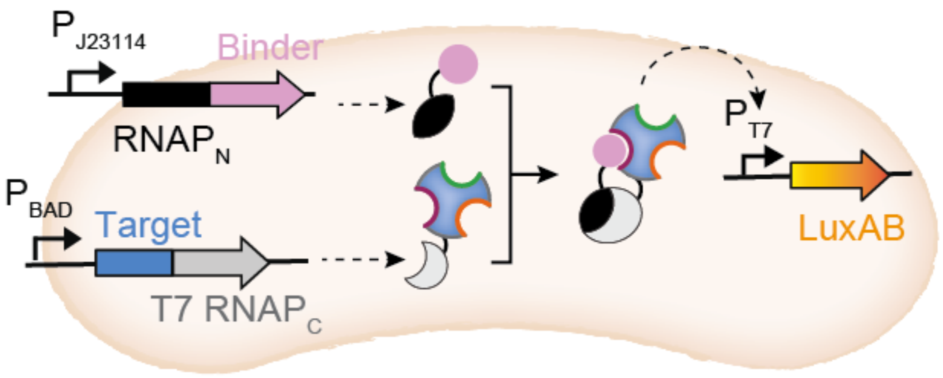


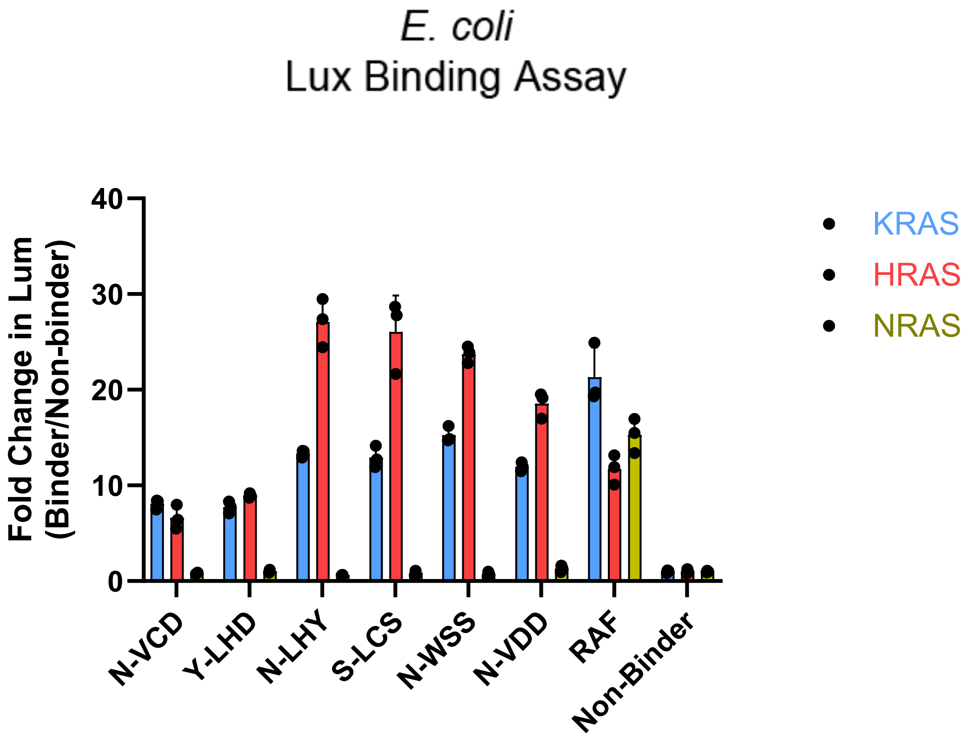


**Supplementary Fig. 1: *E. coli* Binder Lux Assay of Binders Isolated from prior PANCS-Binder Selections.** Schematic of our E. coli split-RNA polymerase luciferase assay to monitor binding between a target and a binding variant. Binders from our prior[1] $10^{10}$ affibody library screen (KRAS G12D vs $ZB_{neg}$) were tested in the *E. coli* luciferase assay for their ability to bind KRAS WT, HRAS, and NRAS (error bars indicate SD, n = 3). Non-binder is the affibody that binds PD-L1 used throughout this work as “Non-binder”.[2] N-VCD is an affibody isolated previously from a separate, smaller ($10^8$) affibody library upon selection for binding to KRAS G12D.[1] RAF indicates the RAS binding domain of RAF (residues 52-121).

| Possible: | 4 | 4,15,6 | 10,15,2 | 15,15 | 8,15 | 8,15 | 3,15 | 15,2 |
|---|---|---|---|---|---|---|---|---|
| 30.6% | N | LHY | IHE | GN | VY | II | VN | DN |
| 17.2% | Y | LHD | IVQ | GN | VF | II | VN | DN |
| 13.5% | S | LCS | YTQ | GN | VY | IV | VR | DN |
| 2.2% | N | VDD | ISQ | GN | VY | II | VA | DD |
| 1.4% | N | WSS | YNE | GN | VF | II | VR | DD |
| N/A | N | VCD | YAQ | GN | VY | IV | IR | DN |
| Motif: | X | XXX | IXX | **GN** | **V**Y | **I**I | **V**X | **D**X |

**Supplementary Fig. 2: Affibody Motif for Binding KRAS.** The top five variants that bind KRAS in the KRAS G12D vs $ZB_{neg}$ selection (all library variants above 1% enrichment in our prior $10^{10}$ affibody library screen).[1] The last sequence (N-VCD) is from a separate 108 affibody library screen on KRAS G12D. Above are listed the number of possible residues in the library randomization for each randomized site. Below is the identified minimal motif for binding; if a residue is consistent across at least 5/6 variants then the amino acid is bolded, if two related amino acids (e.g. Y/F) are present in all of the variants then the more prevalent one is shown (not bolded), all other positions with less convergence are listed as X. We can calculate a frequence of variants with this motif in our library design by multiplying the bold residue possibilities and ½ times the non-bold, but specified possibilities: one of these variants should be expected in every $1.8*10^8$ variants; consistent with our finding of one variant from our $10^8$ library and at least 5 in our $10^{10}$ library.

1 ... 63

**HRAS:** MTEYKLVVVGAGGVGKSALTIQLIQNHFVDEYDPTIEDSYRKQVVIDGETCLLDILDTAGQEE

**KRAS:** MTEYKLVVVGAGGVGKSALTIQLIQNHFVDEYDPTIEDSYRKQVVIDGETCLLDILDTAGQEE

**NRAS:** MTEYKLVVVGAGGVGKSALTIQLIQNHFVDEYDPTIEDSYRKQVVIDGETCLLDILDTAGQEE

64 ... 126

**HRAS:** YSAMRDQYMRTGEGFLCVFAINNTKS**FE**D**IHQ**Y**RE**QI**KR**VKDS**D**DVPMVLVGNKCDLAART**V**E

**KRAS:** YSAMRDQYMRTGEGFLCVFAINNTKSFEDIH**H**YREQIKRVKDS**E**DVPMVLVGNKCDL**PS**RTV**D**

**NRAS:** YSAMRDQYMRTGEGFLCVFAINN**S**KSF**A**DI**NL**YREQIKRVKDSDDVPMVLVGNKCDL**PT**RTV**D**

127 ... 189

**HRAS:** S**RQ**AQ**DL**A**RSY**GIPYIETSAKTRQGVEDAFYTLVREIRQHKLRKLNPPDESGPGCMSCKCVLS

**KRAS:** **TK**QAQDLARSYGIP**F**IETSAKTRQGV**D**DAFYTLVREIR**K**HKEKMSKDGKKKKKKSKTKCVIM

**NRAS:** **TK**QA**HE**LA**K**SYGIP**F**IETSAKTRQGVEDAFYTLVREIRQ**Y**RMKKLNSSDDGTQGCMGLPCVVM

**Supplementary Figure 3. Sequence alignment of HRAS, KRAS, and NRAS Isoforms.** Each RAS isoform (uniprot IDs: P01112, P01116 (isoform 4b), P01111) aligned with mutations from HRAS shown in blue for KRAS and green for NRAS. Bolding and underling in the HRAS sequence indicates residues 4 angstroms or closer to N-LHY in the HRAS:N-LHY structure.

**Supplementary Table 1: Data processing and refinement statistics of NLHY-HRAS**

| | |
|---|---|
| **Data processing** | |
| Wavelength (Å) | 0.9796 |
| Resolution range (Å)[a] | 280.64 – 1.80 (1.83 – 1.80) |
| Space group | $P6_122$ |
| Unit cell parameters (Å) | $a$ = 55.51, $c$ = 280.64 |
| Unique reflections | 25,123 (1,177) |
| Multiplicity | 34.5 (19.9) |
| Completeness (%) | 100 (97.7) |
| <I/σI> | 20.7 (1.4) |
| $R_{merge}$[b] | 0.106 (3.437) |
| CC1/2[c] | (0.543) |
| **Refinement** | |
| Resolution (Å) | 48.09 – 1.80 |
| Reflections work/test set | 23,192/1,738 |
| $R_{work}$/ $R_{free}$[d] | 0.202/0.224 |
| Average B factor ($Å^2$) (No of atoms) | |
| macromolecule | 51.49 (1,632) |
| ligand | 47.98 (32) |
| solvent | 48.90 (87) |
| Rmsd bond lengths (Å) | 0.010 |
| Rmsd bond angles (°) | 1.840 |
| Ramachandran favored[e] (%) | 98.48 |
| Ramachandran outliers | 0 |
| Clashscore[e] | 3.06 |
| PDB ID | 9Z5W |

[a] Values in parentheses correspond to the highest resolution shell.
[b] Rmerge = ΣhΣj|Ihj–<Ih>|/ΣhΣjIhj, where Ihj is the intensity of observation j of reflection h.
[c] As defined by [3]
[d]R = Σh|Fo|–|Fc|/Σh|Fo| for all reflections, where Fo and Fc are observed and calculated structure factors, respectively. Rfree is calculated analogously for the test reflections, randomly selected, and excluded from the refinement.
[e]As defined by Molprobity [4] and implemented in Phenix.

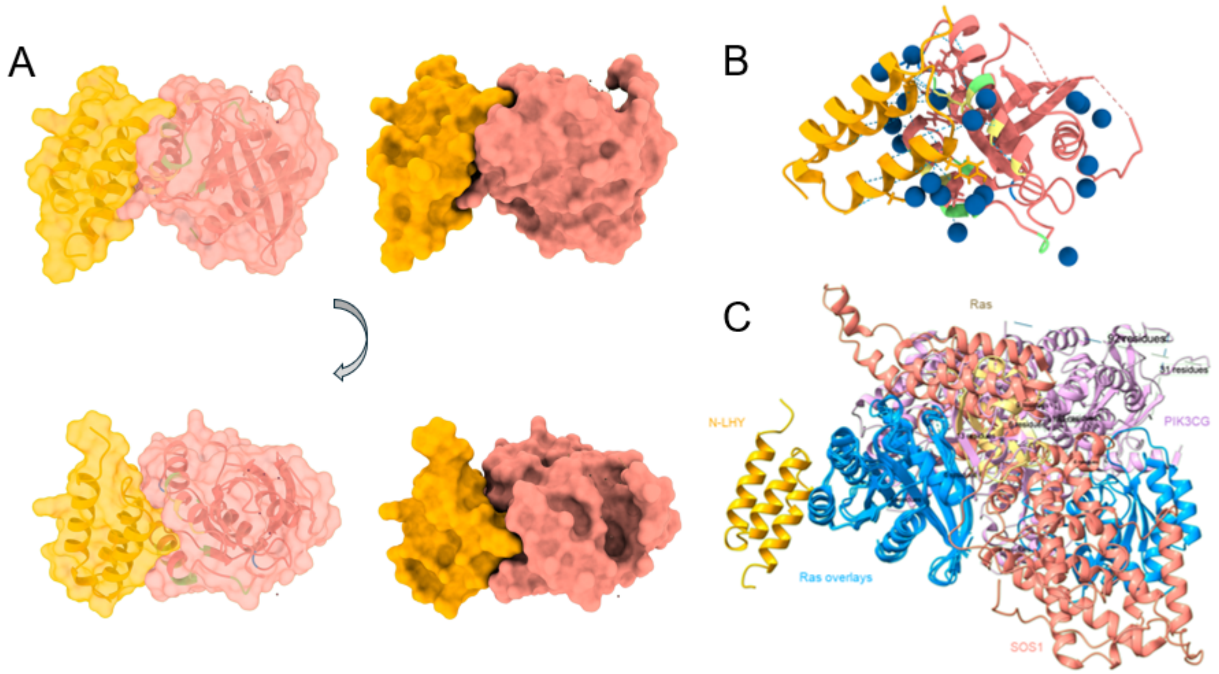


**Supplementary Fig. 4: Additional Details of N-LHY-HRAS Binding Interaction. A)** The N-LHY-HRAS interaction shown in rotated views. **B)** Structured waters were found to bridge several interactions between N-LHY and HRAS that AF3 predicted were direct interactions. **C)** NLHY (orange) binds to an alternative face of RAS (blue) than native PPIs such as PIK3CG (1HE8), SOS1 (1NVW), and RAF (6VJJ).

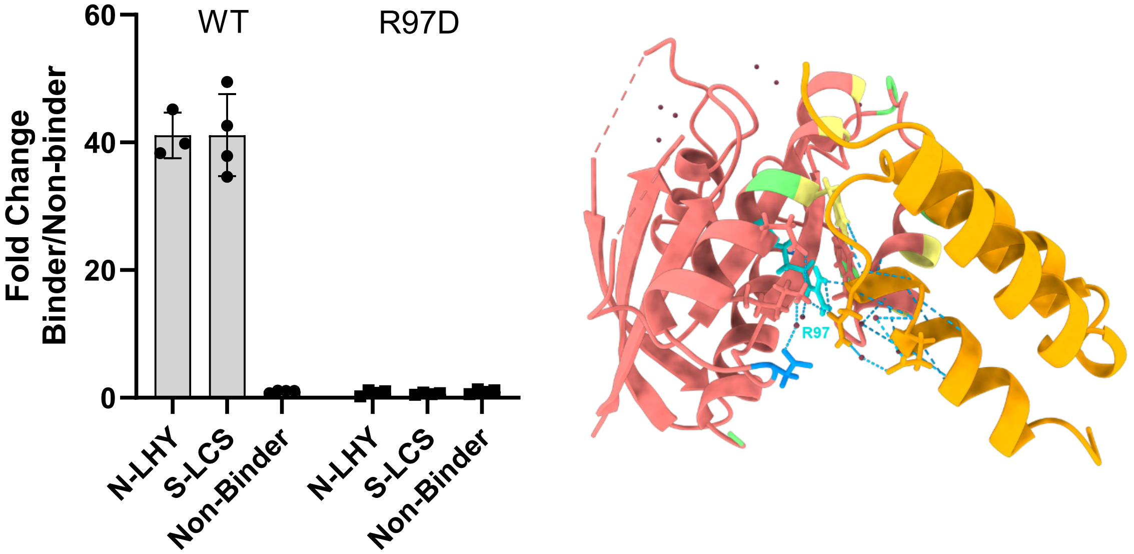


**Supplementary Fig. 5: Mutation that disrupts N-LHY binding to HRAS**

HRAS binders from the affibody screen (HRAS vs $ZB_{neg}$) were tested in the *E. coli* luciferase assay for their ability to bind HRAS R97D (n = 4 replicates). The signal for each target (HRAS or HRAS mutant) is normalized to the non-binding affibody for that target (n=4, error bars indicate SD). In the HRAS-NLHY crystal structure, R97 (indicated in cyan) mediates two contacts with water and two hydrogen bonds (indicted with dashed lines) with N-LHY to the motif residues GN (with the main chain carbonyl and side chain carbonyl, respectively).

| Phage RNAP$_N$ Variant | KRAS vs ZB$_{neg}$ selection strain | HRAS vs ZB$_{neg}$ selection strain | HRAS vs KRAS selection strain | RAF vs KRAS selection strain |
|---|---|---|---|---|
| RAF | Yes | Yes | No | No |
| Affibody (RAF) | No | No | No | Yes |
| N-VCD | Yes | Yes | No | No |
| N-LHY | Yes | Yes | No | No |
| S-LCS | Yes | Yes | No | No |

**Supplementary Figure 6**. **Optimized KRAS –AP expression levels**. Activity-dependent plaque assay results (yes or no) for phage carrying nonspecific RAS binders (Affibody N-VCD, N-LHY, S-LCS or the Ras binding domain of RAF) and an off-target non-binding Affibody that binds RAF[5]. We confirmed that our standard RBS expression levels for our -AP (sd8/sd8) were sufficient to prevent non-specific binder activity dependent plaques, but not too high to prevent a specific binder from replicating. Note that KRAS, HRAS, and RAF have similar expression levels in our system[1] allowing for this comparison. These results suggest that a specific HRAS binder would be able to replicate on the HRAS vs KRAS selection strain.

**Supplementary Table 2. RAS PANCS-spec-Binder Selection Endpoint Titers.** Final titers in Plaque Forming Units/mL (PFU/mL) measured by activity-independent plaque assay for the final passage of each PANCS-binder against RAS isoforms. Screens were as described in the methods, and final titers were collected immediately after the termination of each screen.

| On-Target (+AP) | Off-Target (–AP) | Final Titer (PFU/mL) |
|---|---|---|
| KRAS G12D | Zipper Peptide | $1.42 \times 10^{11}$ |
| HRAS | Zipper Peptide | $1.65 \times 10^{9}$ |
| HRAS | KRAS | $3.50 \times 10^{10}$ |
| NRAS | Zipper Peptide | $4.61 \times 10^{11}$ |
| NRAS | KRAS | $3.17 \times 10^{10}$ |

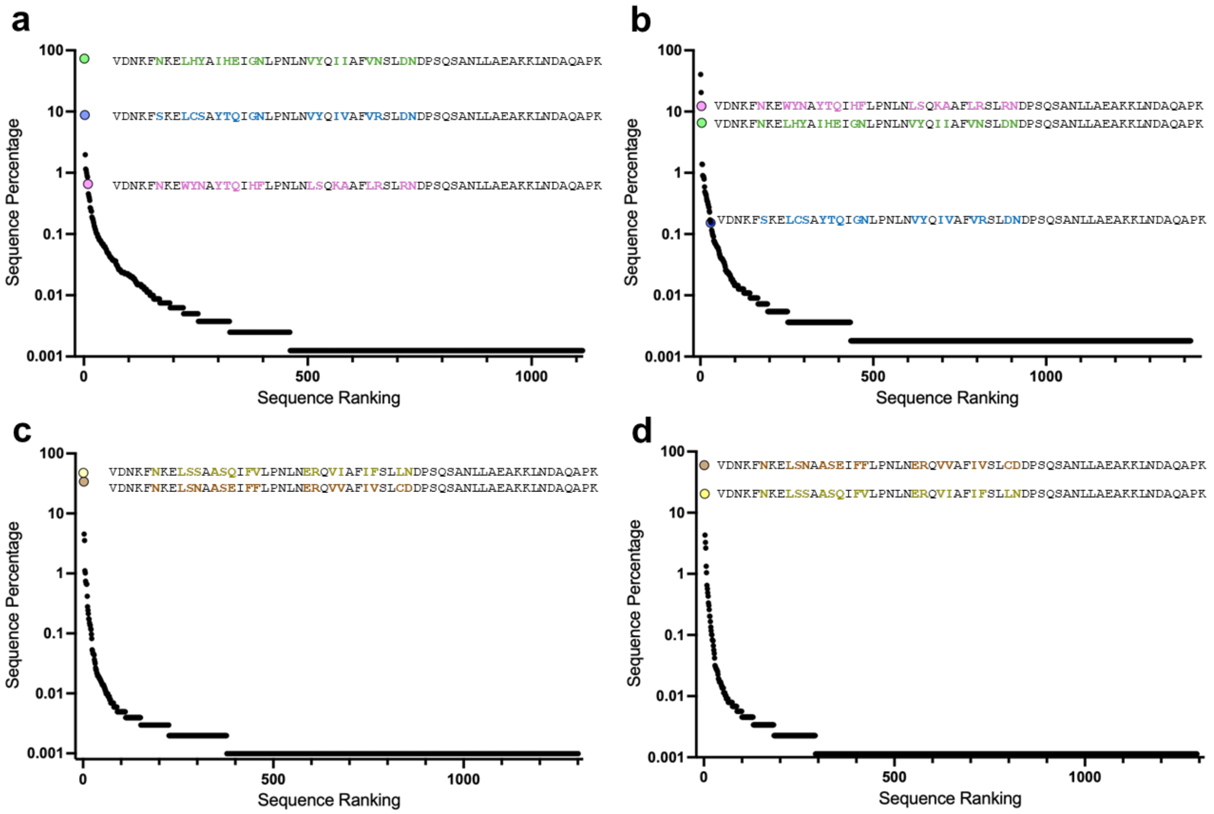


**Supplementary Fig. 7: RAS Selection Endpoint NGS.** Note that KRAS vs $ZB_{neg}$ listed in **Table S2** was previously reported.[1] **a)** Final percentage for all affibody variants enriched in HRAS vs $ZB_{neg}$ selection (80,199 reads, n = 1,124 variants). Variants that were subcloned for further analysis are highlighted (N-LHY: green, S-LCS: blue, N-WYN: pink). **b)** Same as in **a** but for the HRAS vs KRAS selection (55,199 reads, n = 1418 variants). **Fig. 2C** is the direct variant-based comparison of these two screens. **c)** Same as in **a** but for the NRAS vs $ZB_{neg}$ selection (101,123 reads, n = 1,301 variants). Variants that were subcloned for further analysis are highlighted (N-LSS: yellow, N-LSN: brown). **d)** Same as in **c** but for NRAS vs KRAS selection (88,284 reads, n = 1,294 variants).

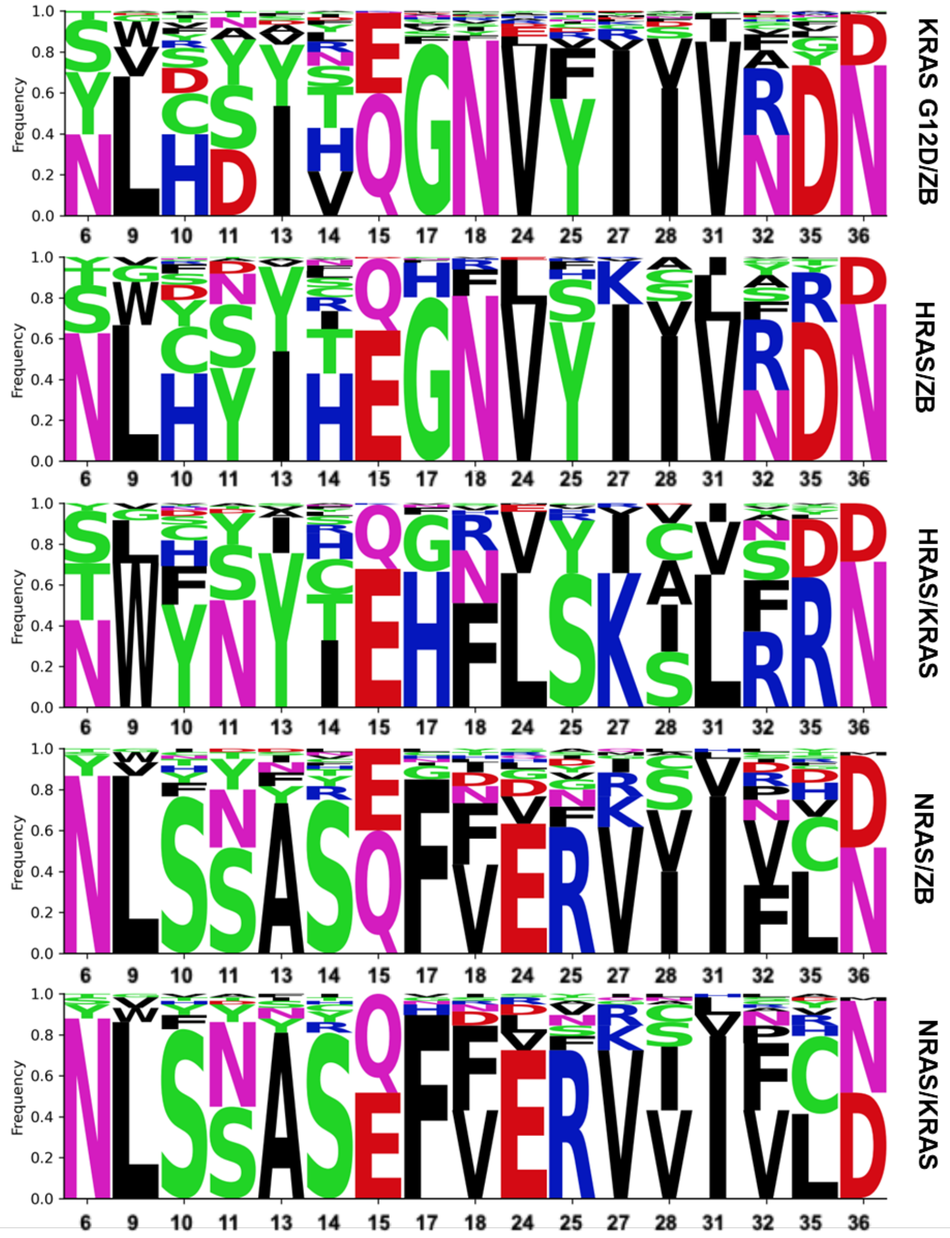


**Supplementary Fig. 8. Sequence logos of selection endpoint variants.** Sequence logos for each RAS targeting screen using $10^{10}$ affibody variant library using 0.01% of the endpoint population as a cut off. Logos are only shown for randomized sites and residues are colored based on biochemical property (black: hydrophobic, green: hydrophilic, blue: positive charge, red: negative charge, pink: polar). Each screen is labeled based on on-target and off-target conditions (i.e. NRAS on-target: KRAS off-target = NRAS/KRAS).

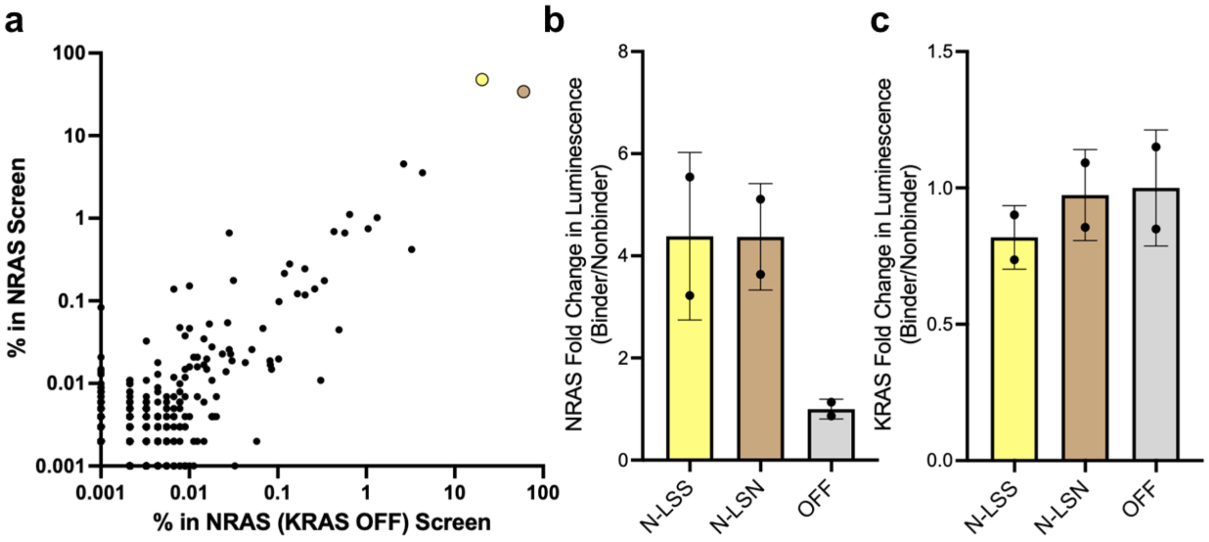


**Supplementary Fig. 9: NRAS Selection Enrichment Comparison and Lux Validation. a,** Sequences for all affibody variants from an initial $10^{10}$ library enriched in screens for NRAS vs KRAS (isoform-selective condition) and NRAS vs $ZB_{neg}$ (non-selective condition) plotted for relative sequence prevalence (n = 2,299). Variants present in only one screen are arbitrarily assigned a prevalence of 0.001% in the opposing screen to facilitate a log-log graph. Correlation analysis shows statistically significant correlation: Pearson r = 0.8116, $R^2$ = 0.6588, $p < 0.0001$ (****). **b,** NRAS binders from the affibody screens (N-LSS: yellow, N-LSN: blue) were tested in the *E. coli* luciferase assay for their ability to bind NRAS (n = 2, error bars indicate SD). The signal is normalized to a non-binding affibody (OFF: gray), and the signal is displayed as a fold change. **c,** Same as in **b** but measuring binding to KRAS WT.

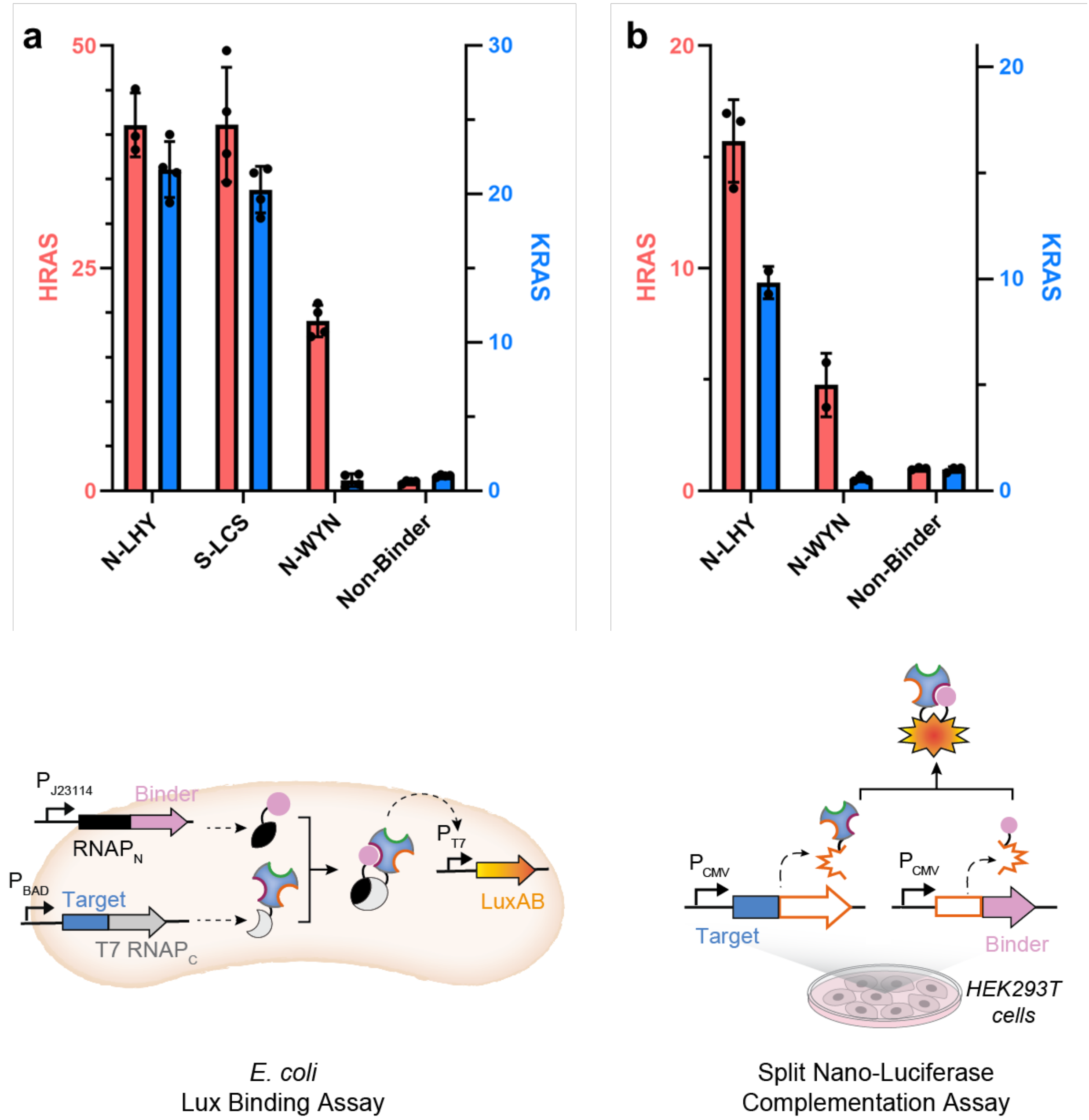


**Supplementary Fig. 10: Lux for 2E and 2F.** **a**, *E. coli* split RNAP luciferase assay for select isolated binders (**Fig. 2E**; n = 4, error bars indicate SD). **b**, Split-luciferase binding assay in HEK293T cells via transfection of RAS isoform fused c-terminally to the c-terminal portion of split-Nano luciferase and binders fused n-terminally to the n-terminal portion of split Nano luciferase (**Fig. 2F**; n = 3, error bars indicate SD). Cartoons indicate the assay design.

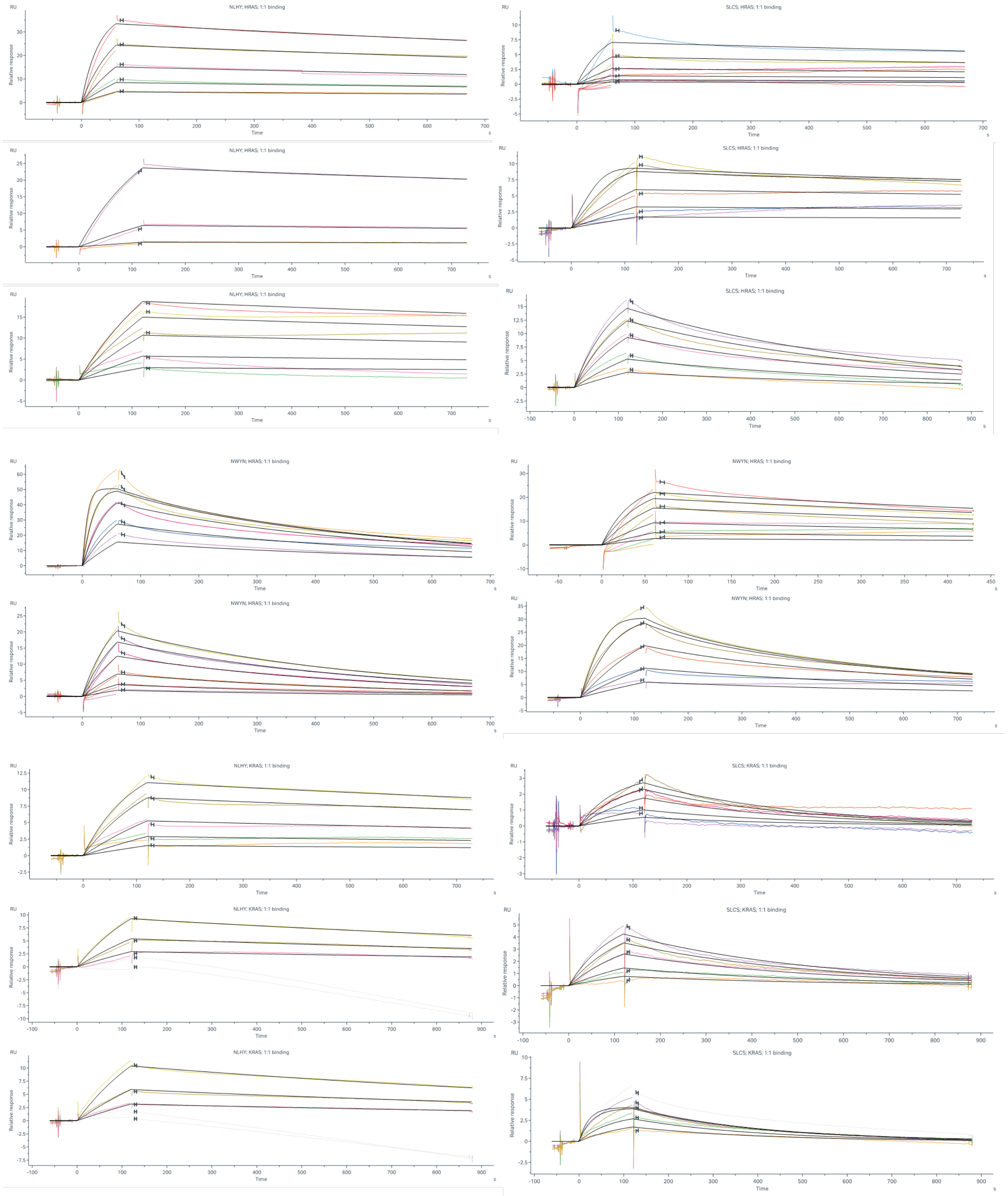


**Supplementary Fig. 11:** Surface Plasmon Resonance (SPR) dose-response and kinetic model fits for Kds shown in Fig. 2D (Supplementary Table 3 for details of model fits). Each target-binder pair required a different concentration range: HRAS with N-LHY (160, 80, 40, 20, 10, 0 nM), with S-LCS (800, 400, 200, 100, 50, 0 nM), and with N-WYN (400, 200, 100, 50, 25, 0 nM); KRAS with N-LHY (80, 40, 20, 10, 5, 0 nM), with S-LCS (800, 400, 200, 100, 50, 0 nM); NRAS with N-LSS and with N-LSN (4000, 3000, 2000, 1000, 500, 0 nM). All interactions were measured in at least duplicate and all data was fit to a 1:1 binding model. (Continues on next page)

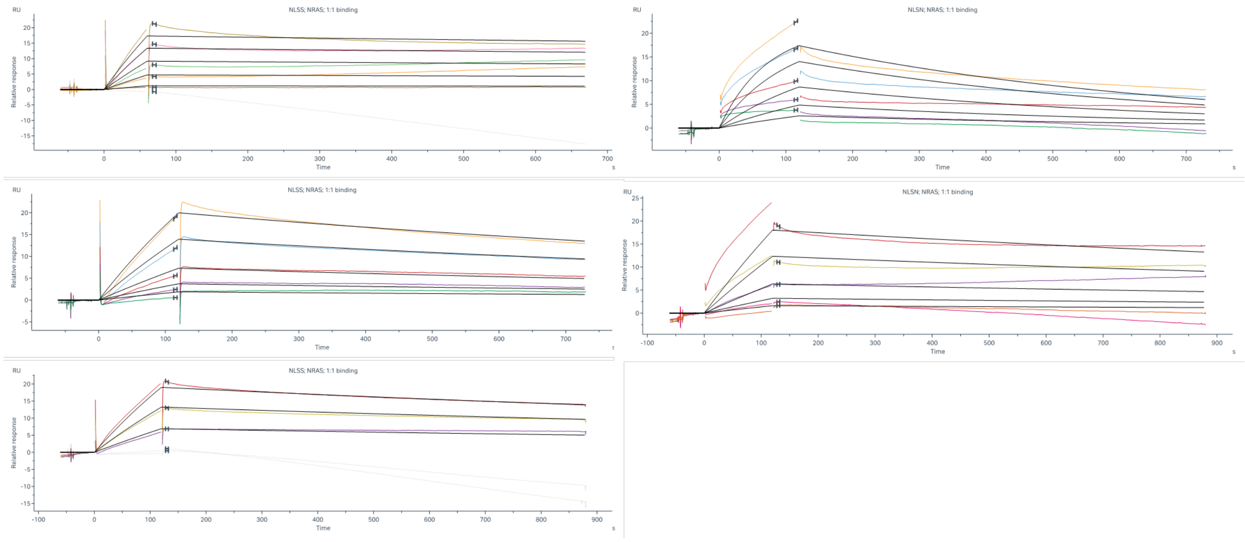


**Supplementary Fig. 11** *(Continued)*

**Supplementary Table 3: SPR Fits.** Kinetic measurements for individual replicates of surface plasmon resonance between a fixed target (RAS) and varied binder concentrations. Average affinity is calculated from the average kinetic parameters. KRAS is KRAS 1-169, HRAS is HRAS 1-166, NRAS is NRAS-166.

| Target | Binder | Replicate | $k_a$ ($M^{-1}s^{-1}$) | $k_d$ ($s^{-1}$) | $K_d$ (nM) |
|---|---|---|---|---|---|
| HRAS | N-LHY | 1 | 8.25E+04 | 3.94E-04 | 4.78 |
| HRAS | N-LHY | 2 | 7.00E+04 | 2.54E-04 | 3.63 |
| HRAS | N-LHY | 3 | 6.33E+04 | 2.81E-04 | 4.44 |
| **HRAS** | **N-LHY** | **Average** | **7.19E+04** | **3.10E-04** | **4.30** |
| HRAS | S-LCS | 1 | 2.62E+04 | 3.78E-04 | 14.4 |
| HRAS | S-LCS | 2 | 1.10E+05 | 4.44E-04 | 4.04 |
| HRAS | S-LCS | 3 | 9.44E+03 | 1.74E-03 | 184 |
| **HRAS** | **S-LCS** | **Average** | **4.85E+04** | **8.54E-04** | **17.6** |
| HRAS | N-WYN | 1 | 4.02E+04 | 2.64E-03 | 65.7 |
| HRAS | N-WYN | 2 | 4.03E+04 | 2.35E-03 | 58.3 |
| HRAS | N-WYN | 3 | 7.85E+04 | 1.00E-03 | 12.7 |
| HRAS | N-WYN | 4 | 1.05E+05 | 3.90E-03 | 37.1 |
| **HRAS** | **N-WYN** | **Average** | **6.60E+04** | **2.47E-03** | **37.46** |
| KRAS | N-LHY | 1 | 1.73E+05 | 3.82E-04 | 2.21 |
| KRAS | N-LHY | 2 | 7.11E+04 | 5.63E-04 | 7.92 |
| KRAS | N-LHY | 3 | 5.96E+04 | 6.73E-04 | 11.3 |
| **KRAS** | **N-LHY** | **Average** | **1.01E+05** | **5.39E-04** | **5.33** |
| KRAS | S-LCS | 1 | 2.77E+04 | 3.58E-03 | 129 |
| KRAS | S-LCS | 2 | 8.25E+03 | 2.39E-03 | 290 |
| KRAS | S-LCS | 3 | 4.23E+04 | 3.49E-03 | 82.5 |
| **KRAS** | **S-LCS** | **Average** | **2.61E+04** | **3.15E-03** | **121** |
| NRAS | N-LSS | 1 | 9.00E+02 | 1.70E-04 | 172 |
| NRAS | N-LSS | 2 | 8.02E+02 | 6.46E-04 | 805 |
| NRAS | N-LSS | 3 | 7.41E+02 | 4.17E-04 | 563 |
| **NRAS** | **N-LSS** | **Average** | **8.14E+02** | **4.11E-04** | **505** |
| NRAS | N-LSN | 1 | 4.19E+03 | 1.74E-03 | 415 |
| NRAS | N-LSN | 2 | 4.64E+02 | 4.06E-04 | 875 |
| **NRAS** | **N-LSN** | **Average** | **2.33E+03** | **1.07E-03** | **461** |

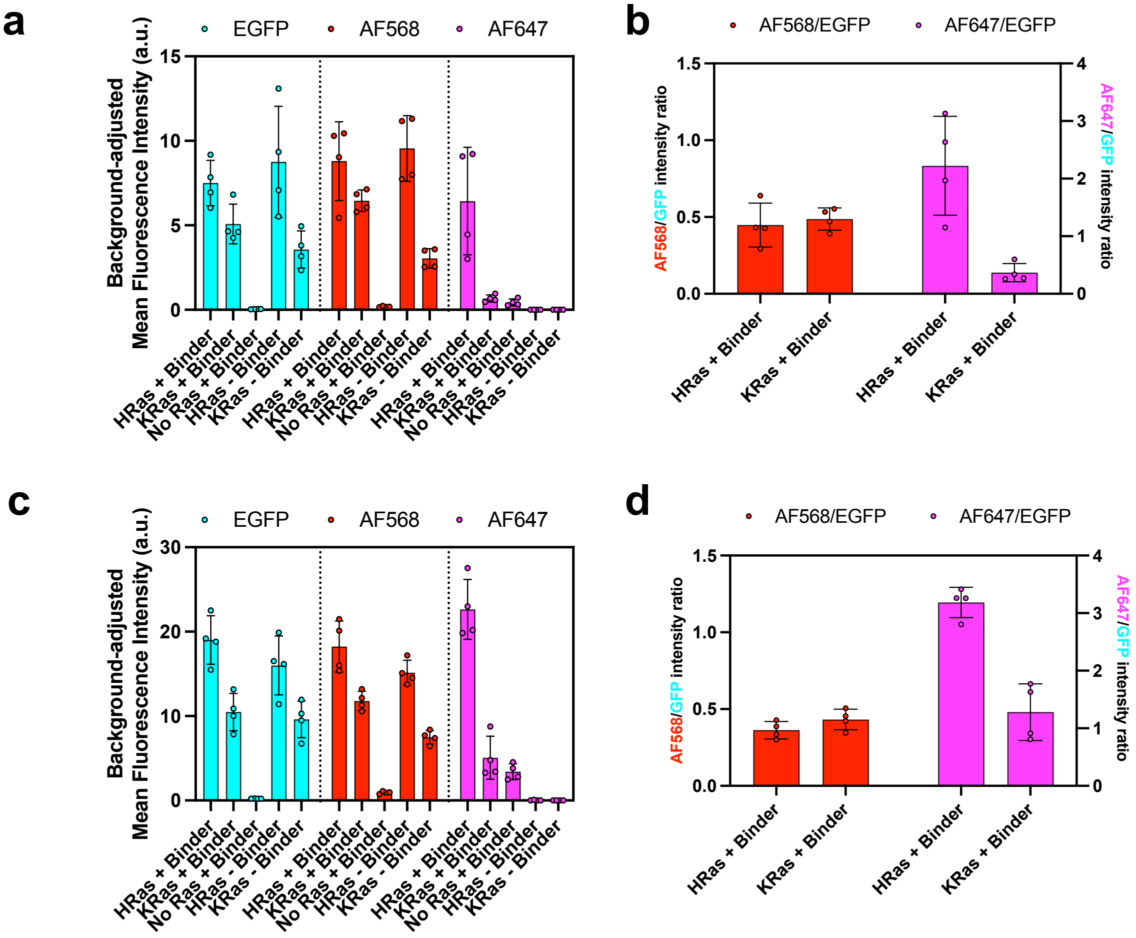


**Supplementary Fig. 12:** Quantification of Fluorescence Intensities showed selective labeling of HRas over KRas using N-WYN binder. **a)** Background-adjusted fluorescence intensity plots of EGFP, AF568, and AF647 from 2.5x zoom-in images **b)** Ratio of AF568 and AF647 over EGFP from 2.5x zoom-in images **c)** Background-adjusted fluorescence intensity plots of EGFP, AF568, and AF647 from 1x zoom-out images **d)** Ratio of AF568 and AF647 over EGFP from 1x zoom-out images. Error bars indicate mean ± s.d.; N = 4.

**See Supplementary Figures S26 and S27 for individual images used in this quantification.**

**Supplementary Table 4: Summary of RAS Specific Binder Variants.** Validated binding highlighted in green for binders with the respective RAS isoform (validated non-binding in red, not tested in grey); see **Supplementary Figure 13** for lux binding assay.

| ID | **Sequence** | **% in HRAS** | **% in KRAS** | **% in NRAS** |
|---|---|---|---|---|
| **KRAS/HRAS Binders** | | | | |
| N-LHY | VDNKFNKELHYAIHEIGNLPNLNVYQIIAFVNSLDNDPSQSANLLAEAKKLNDAQAPK | 73.3 | 30.6 | 0.663 |
| S-LCS | VDNKFSKELCSAYTQIGNLPNLNVYQIVAFVRSLDNDPSQSANLLAEAKKLNDAQAPK | 8.77 | 13.5 | 0.137 |
| N-VDD | VDNKFNKEVDDAISQIGNLPNLNVYQIIAFVASLDDDPSQSANLLAEAKKLNDAQAPK | 0.419 | 2.22 | 0.00989 |
| Y-LHD | VDNKFYKELHDAIVQIGNLPNLNVFQIIAFVNSLDNDPSQSANLLAEAKKLNDAQAPK | 0.0411 | 17.2 | 0.0821 |
| N-WSS | VDNKFNKEWSSAYNEIGNLPNLNVFQIIAFVRSLDDDPSQSANLLAEAKKLNDAQAPK | 0.131 | 1.37 | 0.0129 |
| **HRAS Specific Binders** | | | | |
| T-WYN | VDNKFTKEWYNAYIEIHFLPNLNLSQKSAFLFSLRNDPSQSANLLAEAKKLNDAQAPK | 1.97 | 0.0119 | 0.00989 |
| S-WFS | VDNKFSKEWFSAYCEIHRLPNLNLSQKCAFLSSLRDDPSQSANLLAEAKKLNDAQAPK | 0.998 | 0.001 | 0.001 |
| **HRAS/NRAS Binders** | | | | |
| N-WYN | VDNKFNKEWYNAYTQIHFLPNLNLSQKAAFLRSLRNDPSQSANLLAEAKKLNDAQAPK | 0.657 | 0.00476 | 0.0138 |
| **NRAS Specific Binders** | | | | |
| N-LSS | VDNKFNKELSSAASQIFVLPNLNERQVIAFIFSLLNDPSQSANLLAEAKKLNDAQAPK | 0.190 | 0.276 | 47.8 |
| N-LSN | VDNKFNKELSNAASEIFFLPNLNERQVVAFIVSLCDDPSQSANLLAEAKKLNDAQAPK | 0.101 | 0.233 | 34.1 |
| N-LSS* | VDNKFNKELSSAASQIFVLPNLNERQVVAFIVSLCDDPSQSANLLAEAKKLNDAQAPK | 0.00998 | 0.0238 | 4.54 |

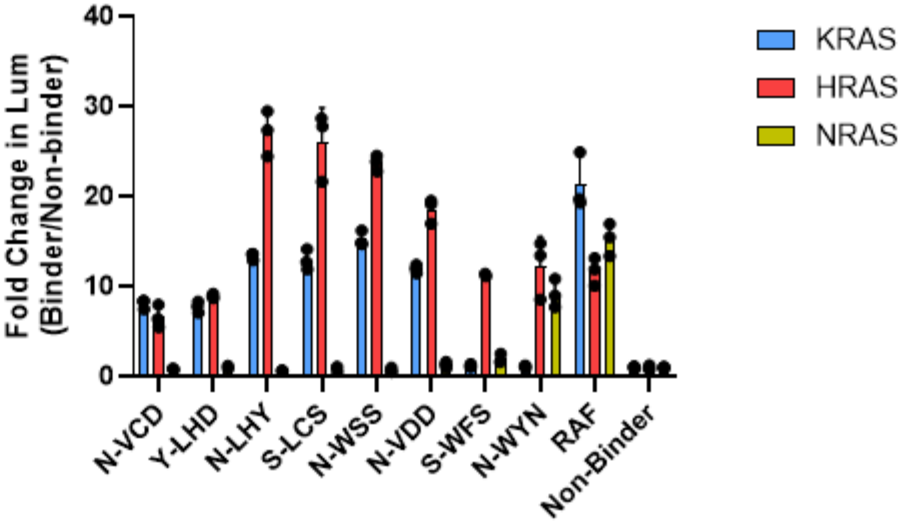


**Supplementary Fig. 13: Cross Reactivity of Isoform Specific RAS Binding Variants.** Binders from our various $10^{10}$ affibody library screens (selective and non-selective vs RAS isoforms, see Supplementary Table 4 for variant details) were tested in the *E. coli* luciferase assay for their ability to bind KRAS WT, HRAS, and NRAS (error bars indicate SD, n = 3). Non-binder is the affibody that binds PD-L1 used throughout this work as "Non-binder".[2] N-VCD is an affibody isolated previously from a separate, smaller ($10^8$) affibody library upon selection for binding to KRAS G12D.[1] RAF indicates the RAS binding domain of RAF (residues 52-121).

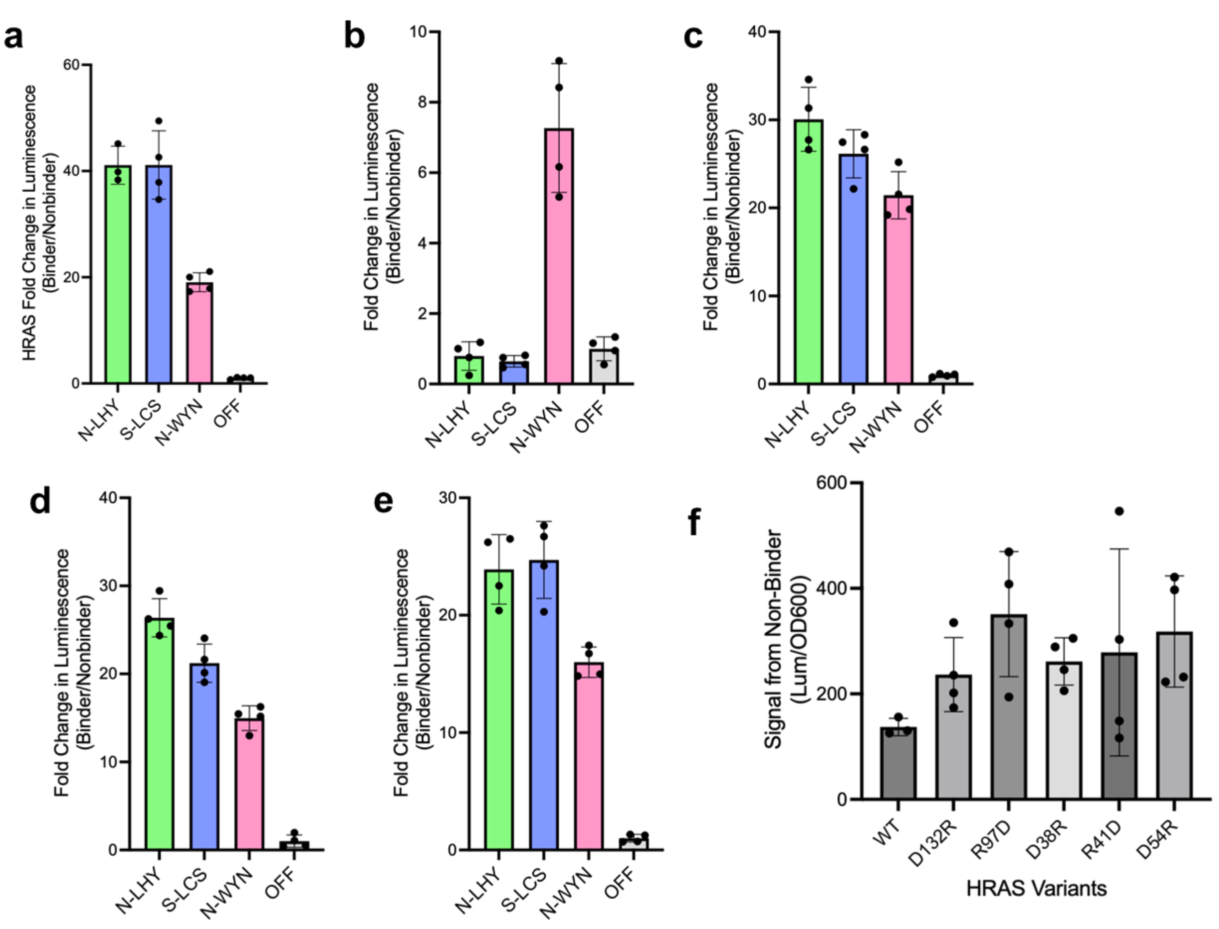


**Supplementary Fig. 14:** Individual replicates for data displayed as a summary in **Fig. 3B. a)** HRAS binders (N-LHY: green, S-LCS: blue, N-WYN: pink) were tested in the *E. coli* luciferase assay for their ability to bind HRAS WT (n = 4, error bars indicate SD). The signal is normalized to a non-binding affibody (OFF: gray), and the signal is displayed as a fold change. **b-e)** Same as in **a,** but for the ability to bind R97D, D38R, R41D, or D54R respectively. **f)** Approximation for the relative expression and stability of HRAS mutants.

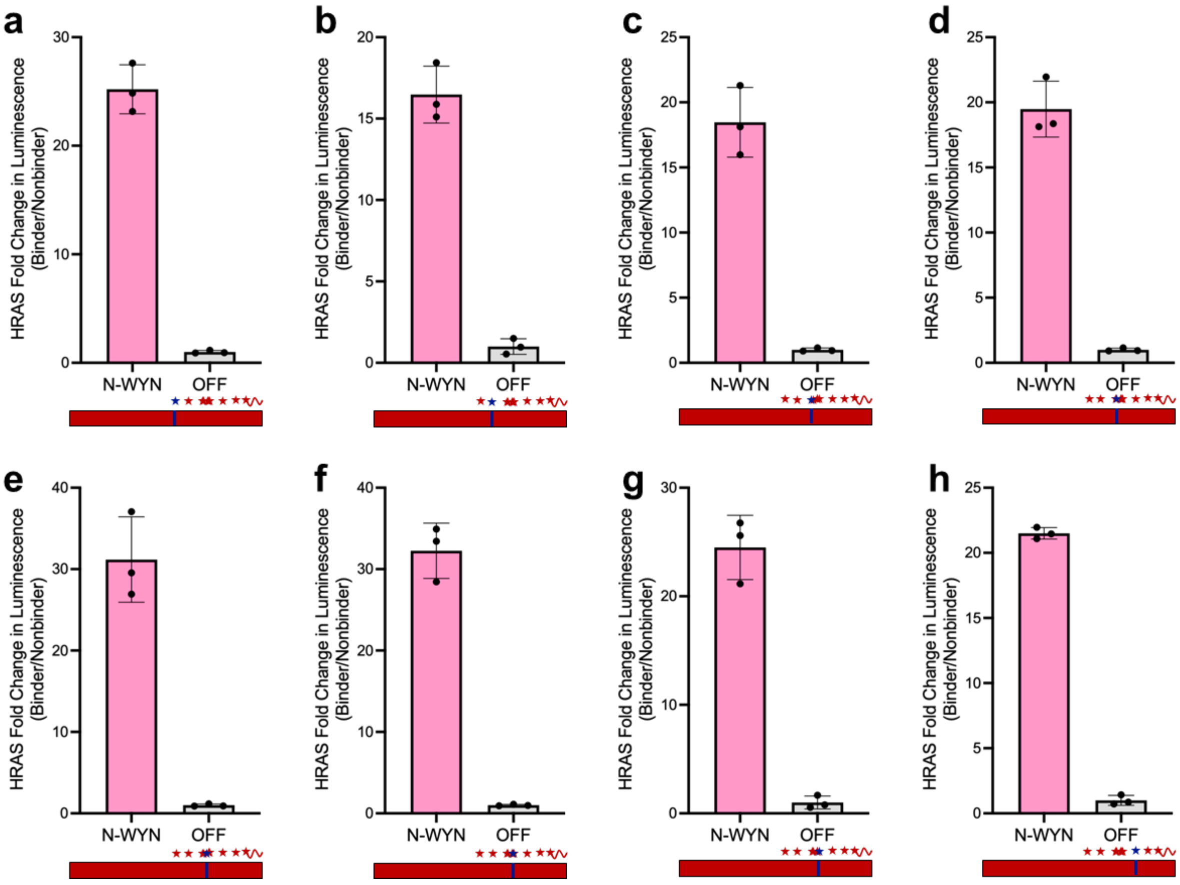


**Supplementary Fig. 15: Lux of Individual Point Mutations to Swap KRAS Mutations into HRAS. a)** HRAS selective binder (N-WYN: pink) was tested in the *E. coli* luciferase assay for its ability to bind HRAS Q95H (n = 3, error bars indicate SD). The signal is normalized to a non-binding affibody (OFF: gray), and the signal is displayed as a fold change. The cartoon below shows the full coding sequence for HRAS (red) with positions that are different between HRAS and KRAS (stars) including the disordered C-terminus (squiggle). Mutations that transition from HRAS (red) to KRAS (blue) are noted. **b-h)** Same as in **a,** but for the ability to bind D107E, A121P, A122S, E126D, S127T, R128K, and Y141F. HRAS D153E was not tested.

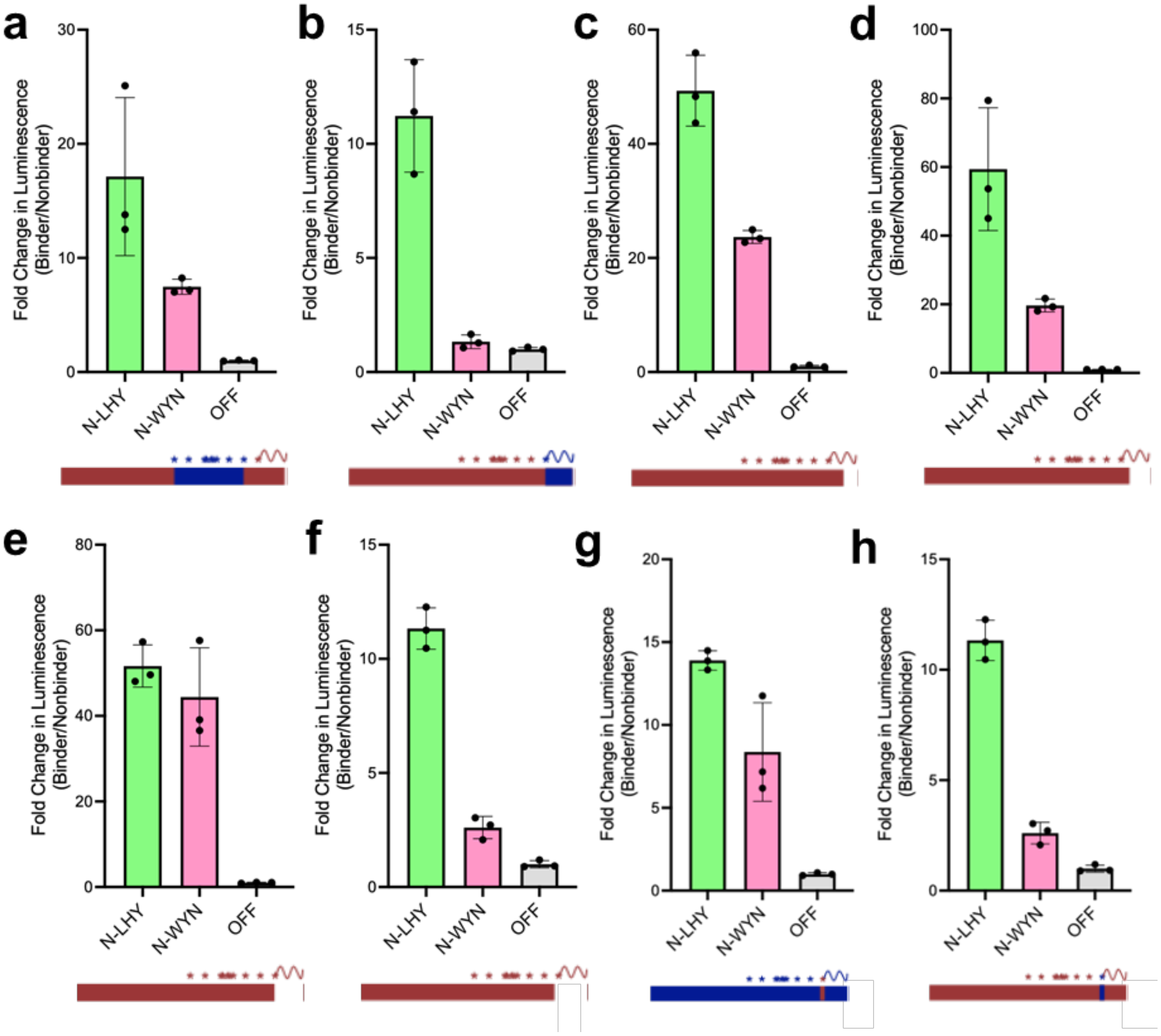


**Supplementary Fig. 16:** Individual replicates for data displayed as a summary in **Fig. 3C-D. a)** HRAS binders (N-LHY: green, N-WYN: pink) were tested in the *E. coli* luciferase assay for the ability to bind HRAS (KRAS 95-153) (n = 3, error bars indicate SD). The signal is normalized to a non-binding affibody (OFF: gray), and the signal is displayed as a fold change. The cartoon below shows the full coding sequence for HRAS (red) with positions that are different between HRAS and KRAS (stars) including the disordered C-terminus (squiggle). Mutations that transition from HRAS (red) to KRAS (blue) are noted. **b-h)** Same as in **a,** but for the ability to bind (**b**) HRAS (KRAS 165-188); HRAS truncations (**c-f**): 1-178, 1-172, 1-166, 1-162; and inversions at position 165: KRAS K165Q (**g**) and HRAS Q165K (**h**).

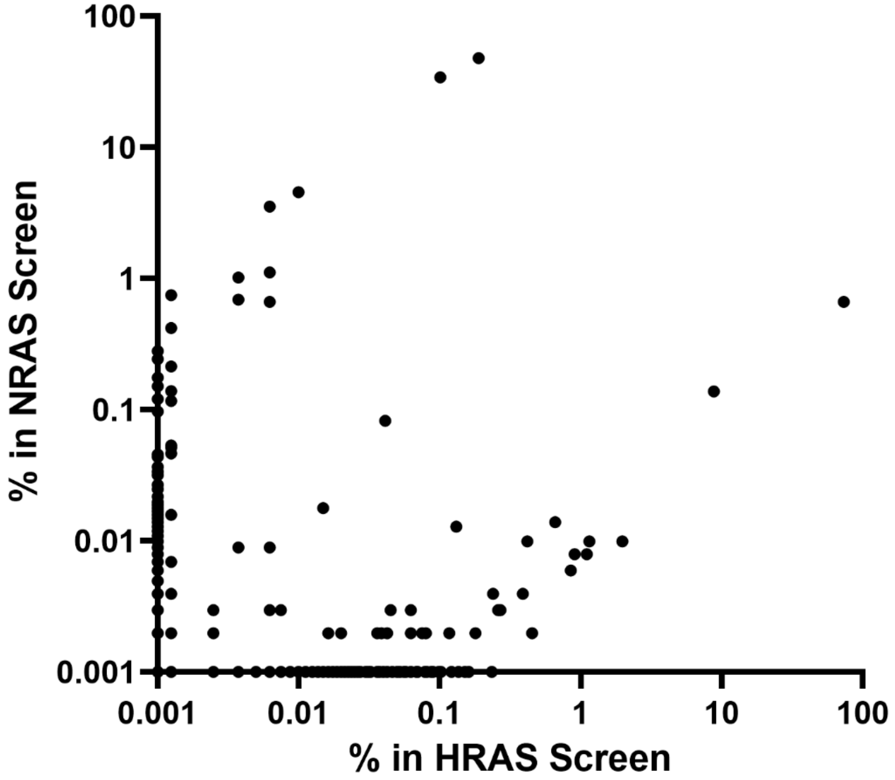


**Supplementary Fig. 17: Comparison of HRAS vs $ZB_{neg}$ and NRAS vs $ZB_{neg}$.** Sequences for all affibody variants from an initial $10^{10}$ library enriched in screens for HRAS vs $ZB_{neg}$ and NRAS vs $ZB_{neg}$ plotted for relative sequence prevalence (n = 7,619). Variants present in only one screen are arbitrarily assigned a prevalence of 0.001% in the opposing screen to facilitate a log-log graph. Correlation analysis shows statistically significant correlation: Pearson r = 0.01398, p = 0.2224 (ns).

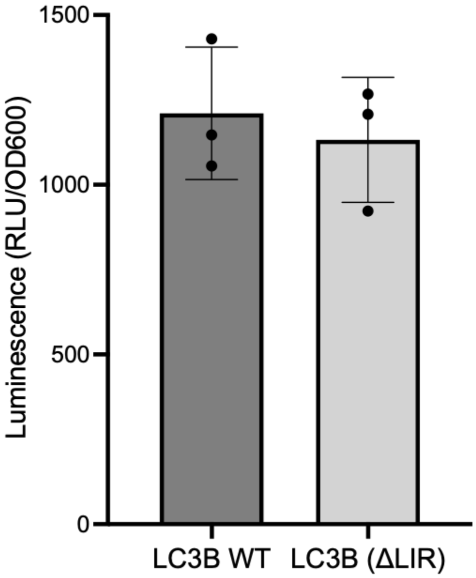


**Supplementary Fig. 18: Expression Level of LC3B WT and LC3B LIR Mutant.** Relative expression of LC3B wild type and the LIR-mutants variant chosen for the -AP based on background luminescence from a non-binding affibody (OFF) (n = 3, error bars indicate SD).

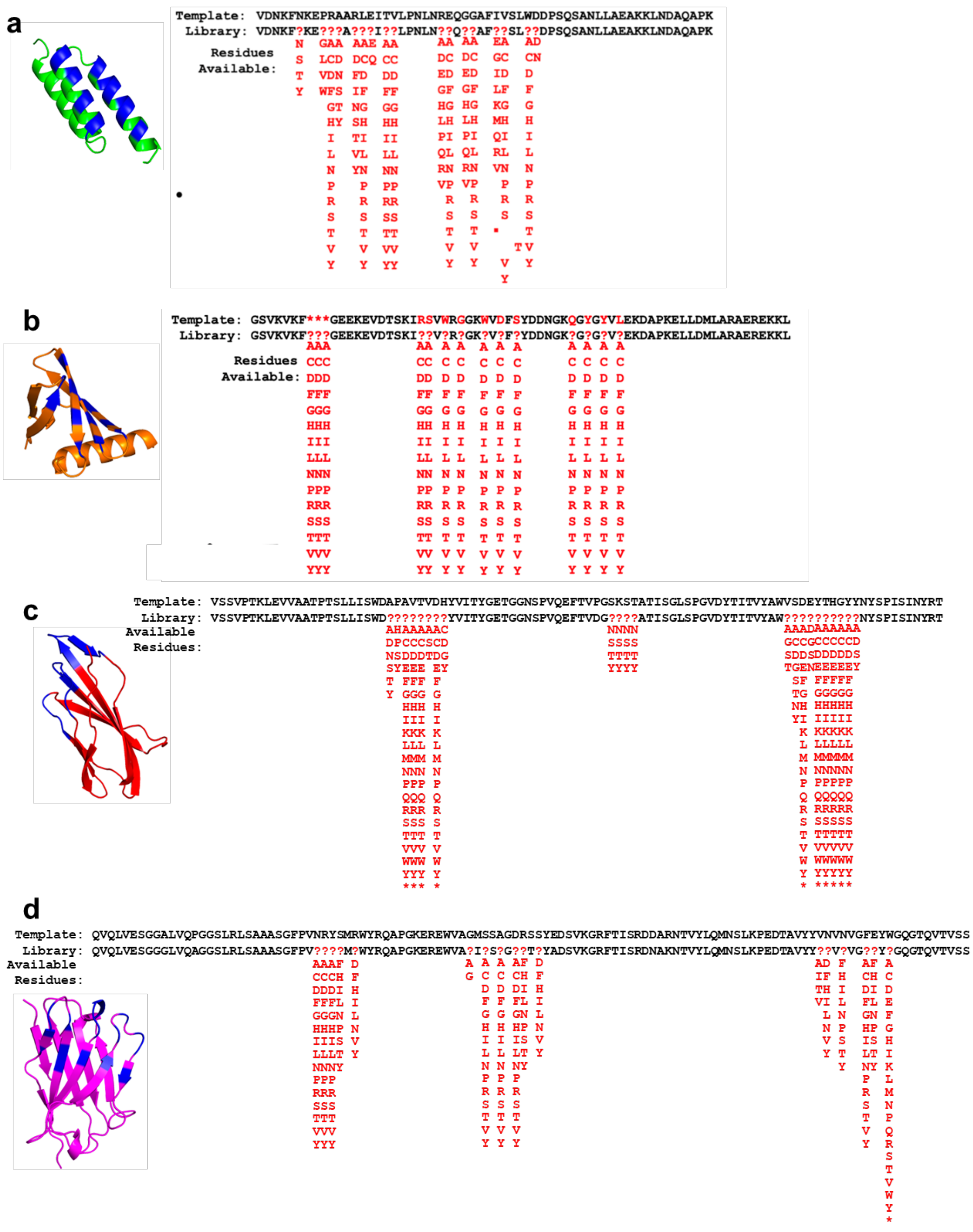


**Supplementary Fig. 19: Mixed Scaffold Library Summary.** Our pooled library consisted of four libraries with designs shown for **a**) affibody, **b**) affitin, **c**) monobody, and **d**) nanobody. Randomized positions indicated in red in the sequences with the possible amino acids based on codon degeneracy listed below. Structures show scaffold with randomized positions indicated in blue. The affibody and affitin libraries were our previously reported ~108 variant libraries. The nanobody and monobody libraries were slightly smaller (as measured by titer 1 hr post transformation), ~5*10^7 and 1*10^7, respectively.

**Supplementary Table 5: LC3B Selection Endpoint Titers**. Final titers in Plaque Forming Units/mL (PFU/mL) measured by activity-independent plaque assay for the final passage of each LC3B PANCS-Binder selection.

| On-Target (+AP) | Off-Target (–AP) | Final Titer (PFU/mL) |
|---|---|---|
| LC3B | Zipper Peptide | $2.38 \times 10^{10}$ |
| LC3B | LC3B | $8.00 \times 10^{3}$ |
| LC3B | LC3B (ΔLIR) | $1.07 \times 10^{10}$ |

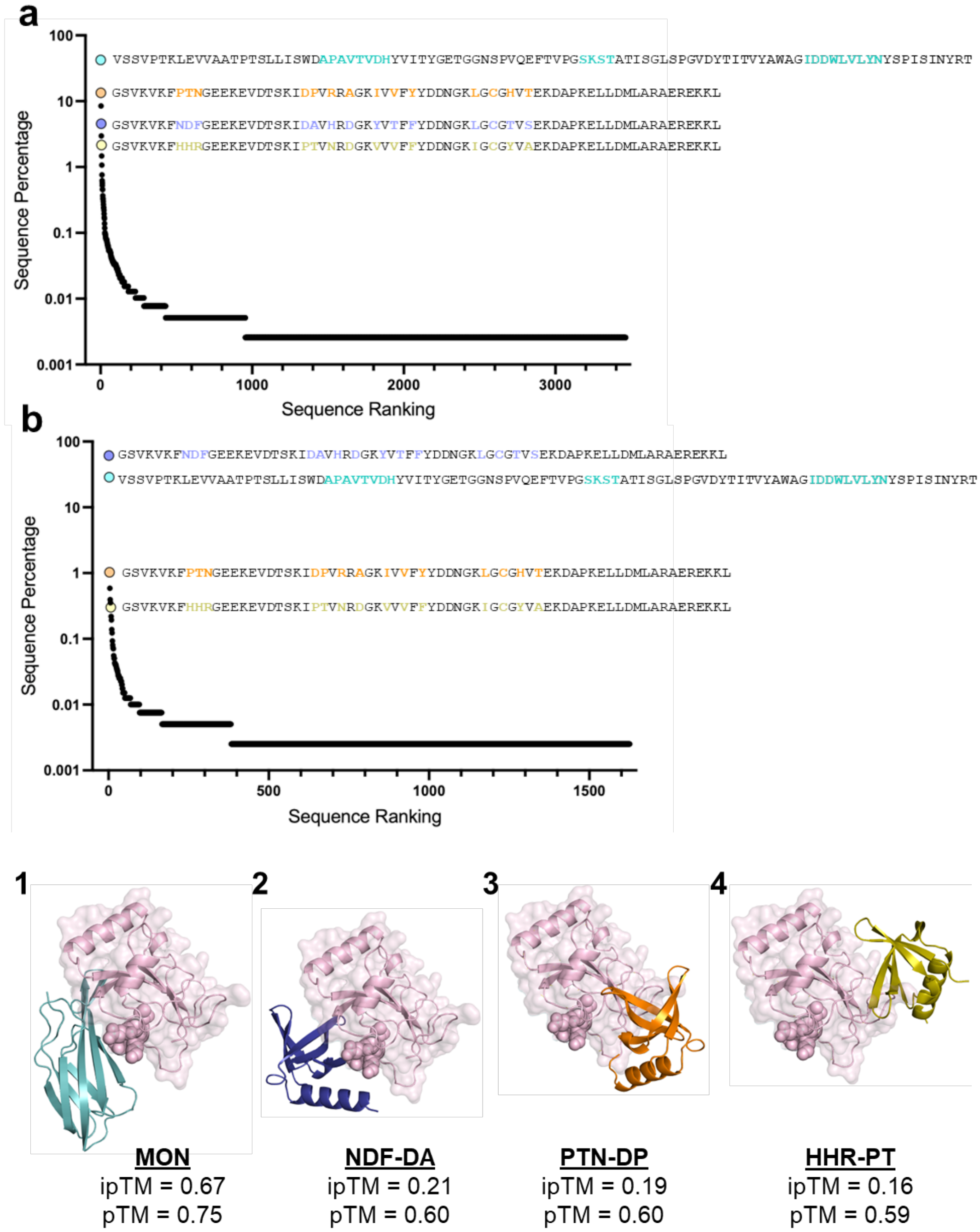


**Supplementary Fig. 20: NGS of LC3B Selections**. **a)** Final percentage for all variants enriched in the LC3B vs $ZB_{neg}$ selection (38,932 reads, n = 3,463 variants). Variants that were subcloned for further analysis are highlighted (MON: teal (**1**), NDF: blue (**2**), PTN: orange (**3**), HHR: yellow (**4**)). **b)** Same as in **S21A** but for the LC3B-targeting screen with epitope-selective conditions (LC3B (ΔLIR)) (39,735 reads, n = 1,628 variants). **Fig. 4E** is the direct variant-based comparison of these two screens. Below are the AlphaFold3 predicted binding complexes of our four isolated binders with LC3B. Residues that were mutated for the LIR-directed screen are shown as balls (E62, K65, I67, R69) and Alphafold3 confidence metrics are listed below (pTM = predicted template modeling score, ipTM = interaction predicted template modeling score).

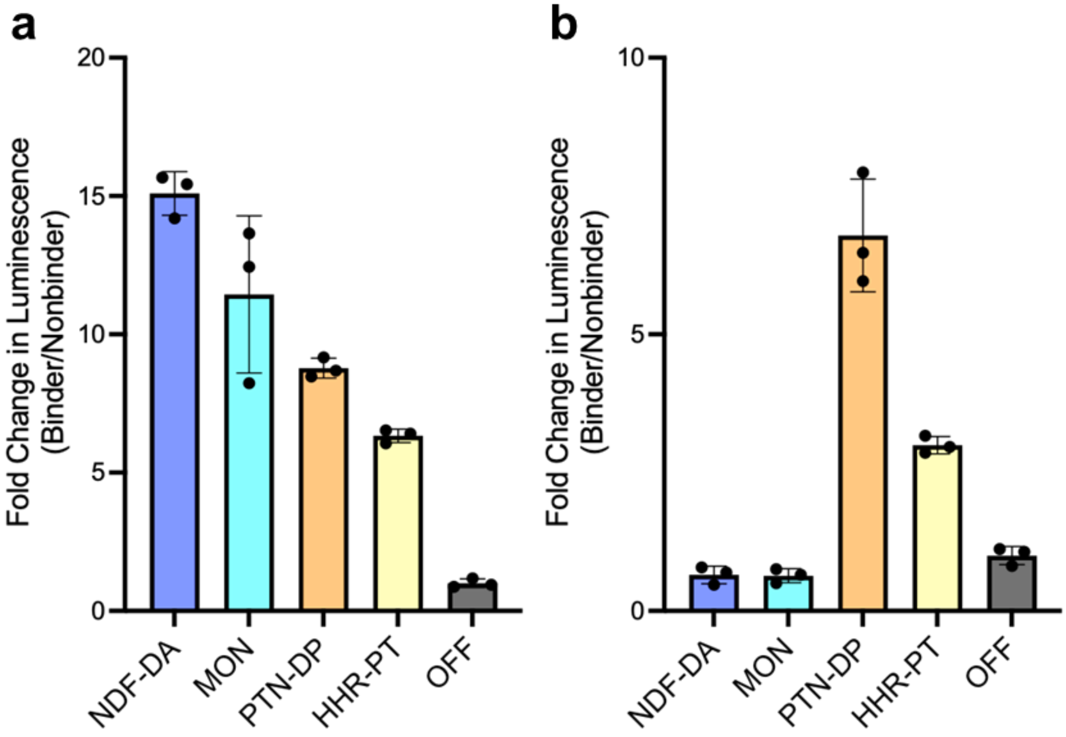


**Supplementary Fig. 21: Lux for 4E Individual Replicates.** Individual replicates for data displayed as a heatmap in **Fig. 4E**. **a**) LC3B binders from the mixed library screen actively enriched with epitope-selective conditions (NDF-DA (**2**): blue, MON: teal (**1**)) or actively de-enriched under epitope-selective conditions (PTN-DP: orange (**3**), HHR-PT: yellow (**4**)) were tested in the *E. coli* luciferase assay for their ability to bind LC3B WT (error bars indicate SD, n = 3). The signal is normalized to our non-binding affibody as an off-target (OFF: gray), and the signal is displayed as a fold change. **b**) Same as in **a** but measuring binding to LC3B with mutations at the LIR face (LC3B (**ΔLIR**)).

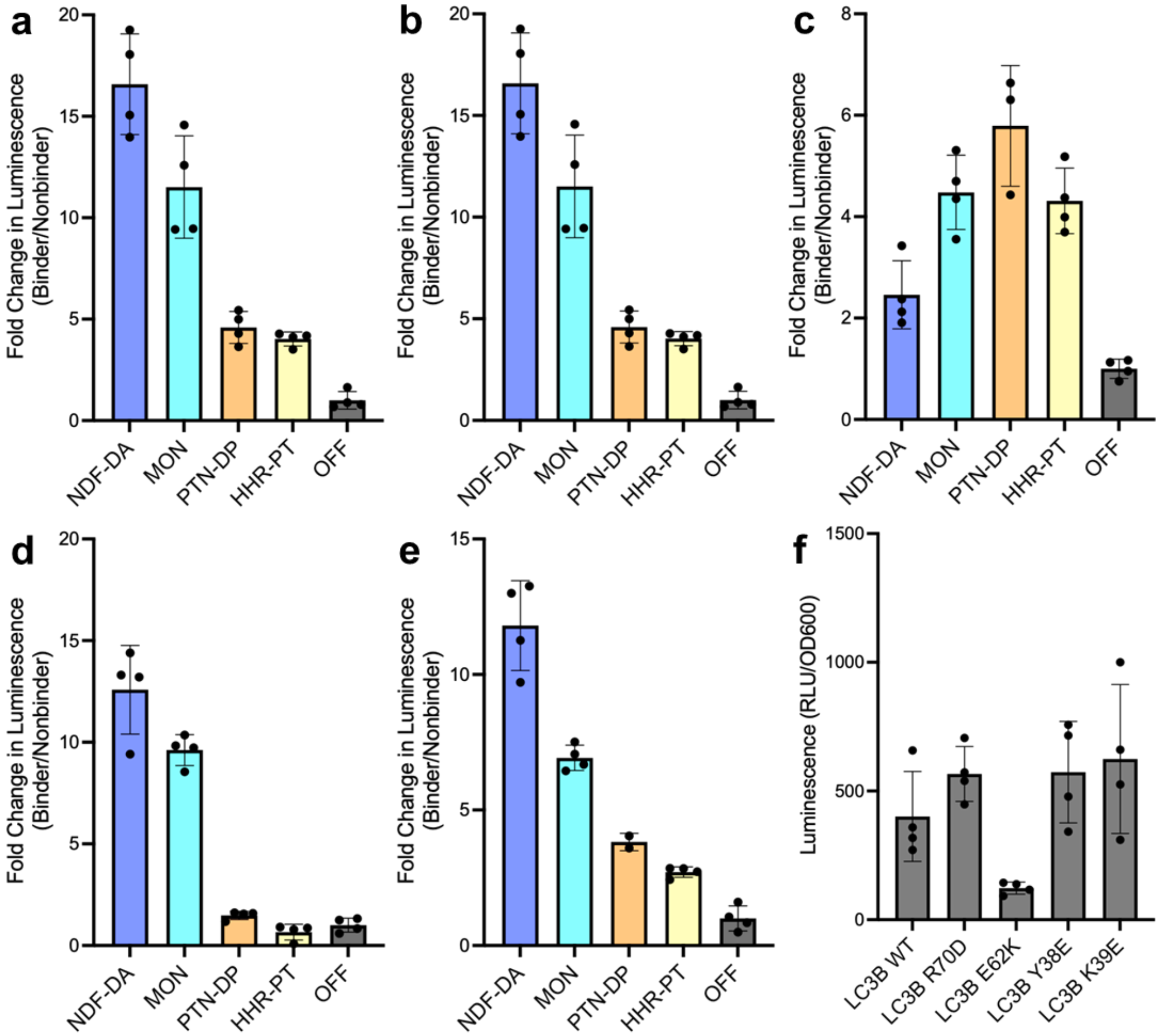


**Supplementary Fig. 22: Lux for 4G Individual Replicates.** Individual replicates for data displayed as a heatmap in **Fig. 4G**. **a**) LC3B binders from the mixed library screen actively enriched with epitope-selective conditions (NDF-DA (**2**): blue, MON: teal (**1**)) or actively de-enriched under epitope-selective conditions (PTN-DP: orange (**3**), HHR-PT: yellow (**4**)) were tested in the *E. coli* luciferase assay for their ability to bind LC3B WT (error bars indicate SD, n = 4). The signal is normalized to our non-binding affibody as an off-target (OFF: gray), and the signal is displayed as a fold change. **b-c**) Same as in **a** but measuring binding to LC3B with mutations at predicted LIR interface (R70D and E62K, respectively). **d-e**) Same as in **a** but measuring binding to LC3B with mutations at predicted secondary interface (Y38E and K39E, respectively). **f**) Relative expression of LC3B mutants based on luminescence from our non-binding affibody as an off-target (error bars indicate SD, n =4).

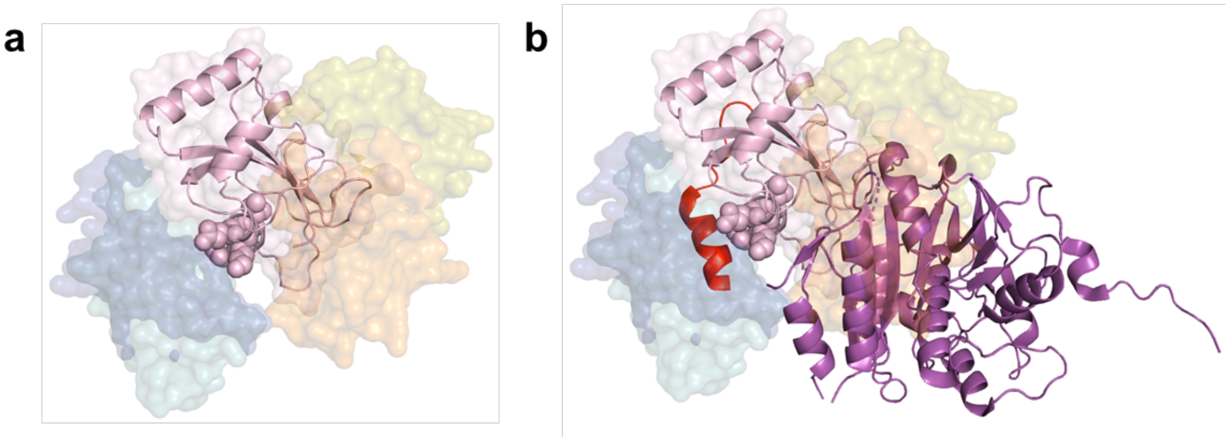

**Supplementary Fig. 23: LC3B:ATG4b Interaction Site**. Comparison binding epitopes of native and our discovered binders. **a**) Overlay of all tested LC3B binders with LC3B (pink) predicted by AlphaFold3 (MON: teal (**1**), NDF: blue (**2**), PTN: orange (**3**), HHR: yellow (**4**); see **Supplemental Figure 20** for individual predictions). Residues that were mutated for the LIR-directed screen are shown as balls (E62, K65, I67, R69). **b**) Same as **a** but including a previously crystallized LIR peptide in complex with LC3B (red, PDB: 5WRD) and a previously crystallized fragment of ATG4B in complex with LC3B (purple, PDB: 2Z0D).

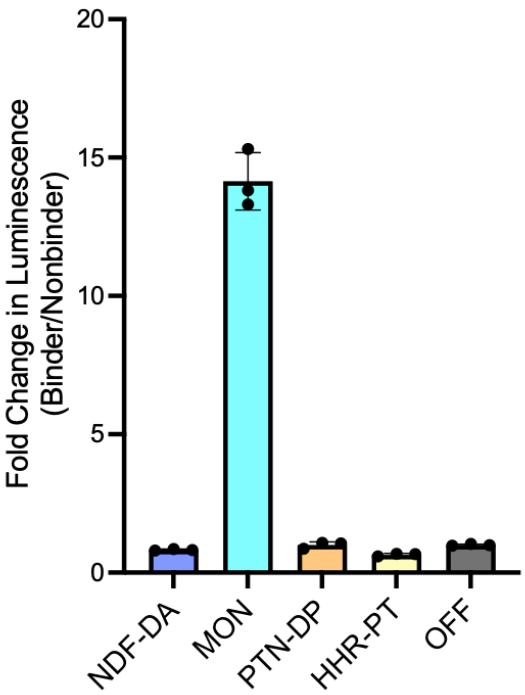


**Supplementary Fig. 24: Lux for 4H Individual Replicates.** Individual replicates for data displayed as a heatmap in **Fig. 4H**. LC3B data shown in **Supplementary Fig. 21**. Binders from the mixed library screen actively enriched with epitope-selective conditions (NDF-DA (**2**): blue, MON: teal (**1**)) or actively de-enriched under epitope-selective conditions (PTN-DP: orange (**3**), HHR-PT: yellow (**4**)) were tested in the *E. coli* luciferase assay for their ability to bind GABARAP (error bars indicate SD, n = 3). The signal is normalized to our non-binding affibody as an off-target (OFF: gray), and the signal is displayed as a fold change.

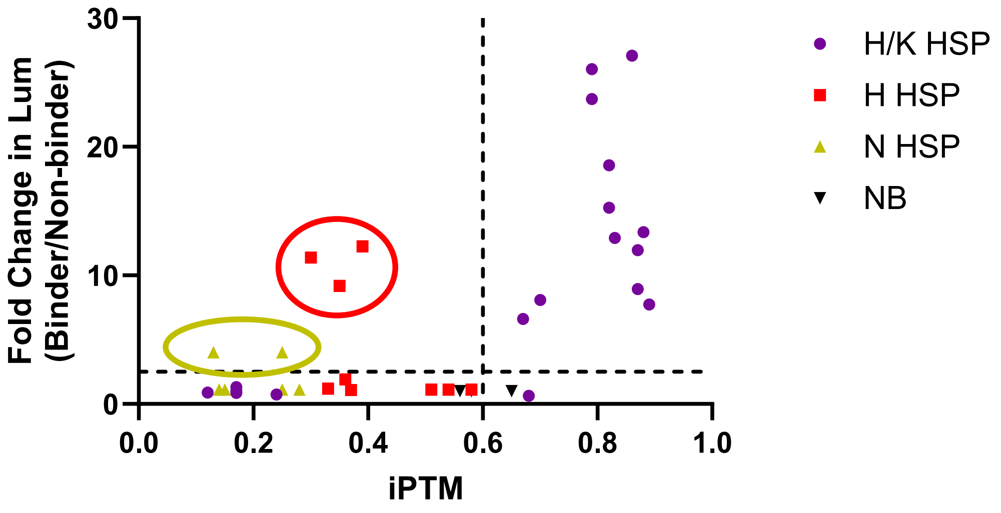


**Supplementary Fig. 25:** AlphaFold3 predicted structures (x-axis has iPTM values) compared to the measured binding in our E. coli luciferase assay (y-axis has fold-change in lux) for RAS binders and RAS isoforms (in **Supplementary Table 4**). The H/K HSP (HRAS/KRAS hot-spot; purple) points refer to variants with the motif (**Supplementary Figure 2**) for binding HRAS and KRAS. For this hotspot, AlphaFold3 does reasonably well with all HRAS and KRAS predictions being >0.6 iPTM (and they all bind) and only one NRAS prediction being >0.6 iPTM (none bind). However, for the other two hotspots (for the HRAS specific motif exemplified by SWFS (red) and for the NRAS specific motif exemplified by NLSS (yellow)), AlphaFold3 fails to predict any binding interactions; additionally, the non-binding (NB, black) affibody is predicted to bind to KRAS and is near the 0.6 threshold for both HRAS and NRAS. These results suggest that AlphaFold3 performance is strongly bifurcated; when it predicts a binding interaction well, it can do so with many variants of the underlying binding motif; however, it still failed to predict binding for the majority of our motifs.

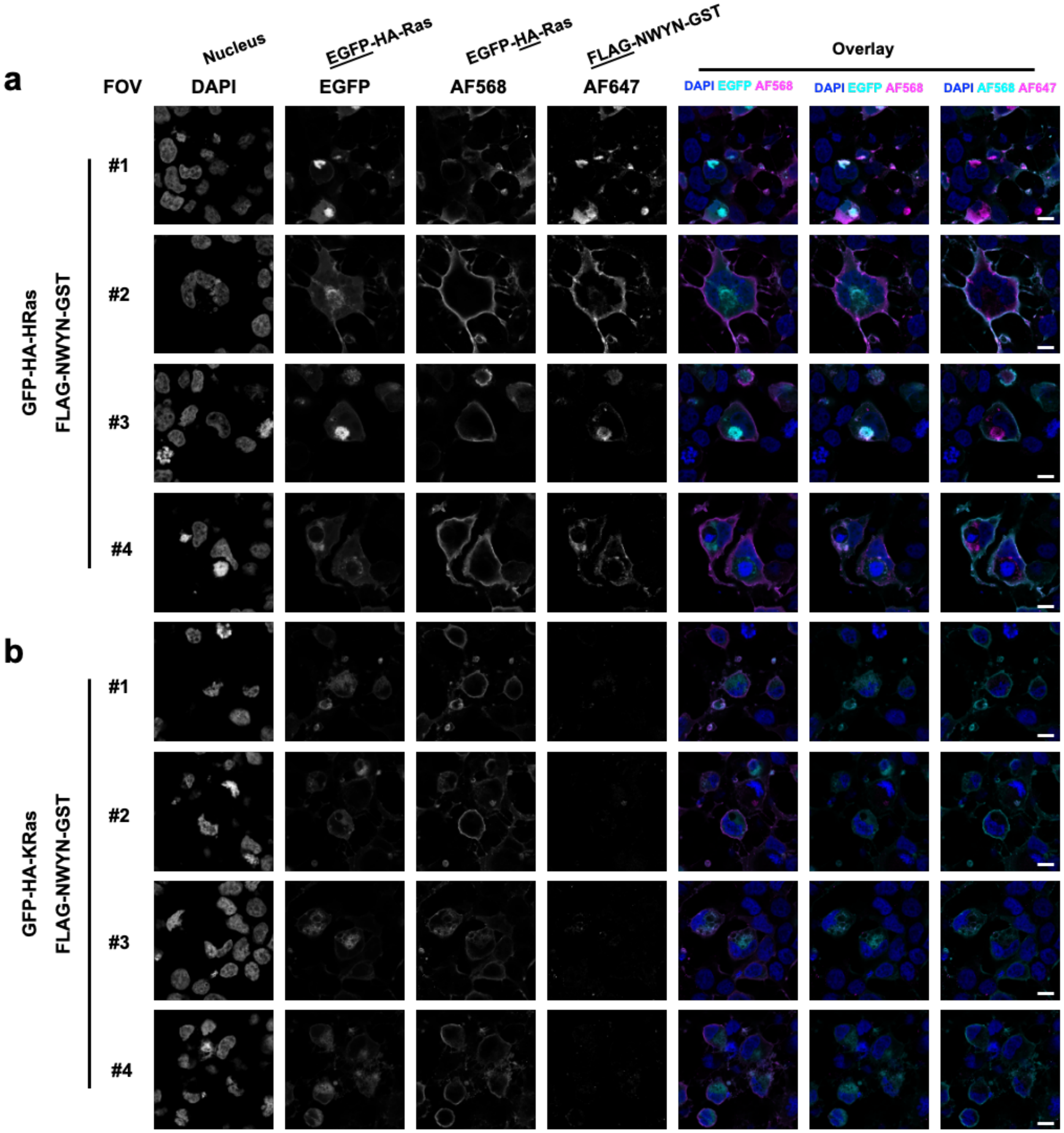


Cont.

**Supplementary Fig. 26: Confocal microscopy demonstrated HRas-selective immunofluorescence staining using purified recombinant N-WYN binder.** HEK293T cells overexpressing GFP-HA-HRas or GFP-HA-KRas fusion protein were stained anti-HA/anti-rabbit-AF568, 3xFLAG-GST-NWYN/anti-FLAG/anti-Ms-AF647, and DAPI **(a,b)**. Untransfected and binder-omitted controls were included as background references. **(c-d)**. Four fields of views (FOVs) were shown per condition. *Top insets and underlined labels denote the intended viewing targets and corresponding color overlay schemes. Images were acquired using a 63x objective lens with 2.5x confocal zoom. Scale Bar, 10 μm. (Continues on next page)*

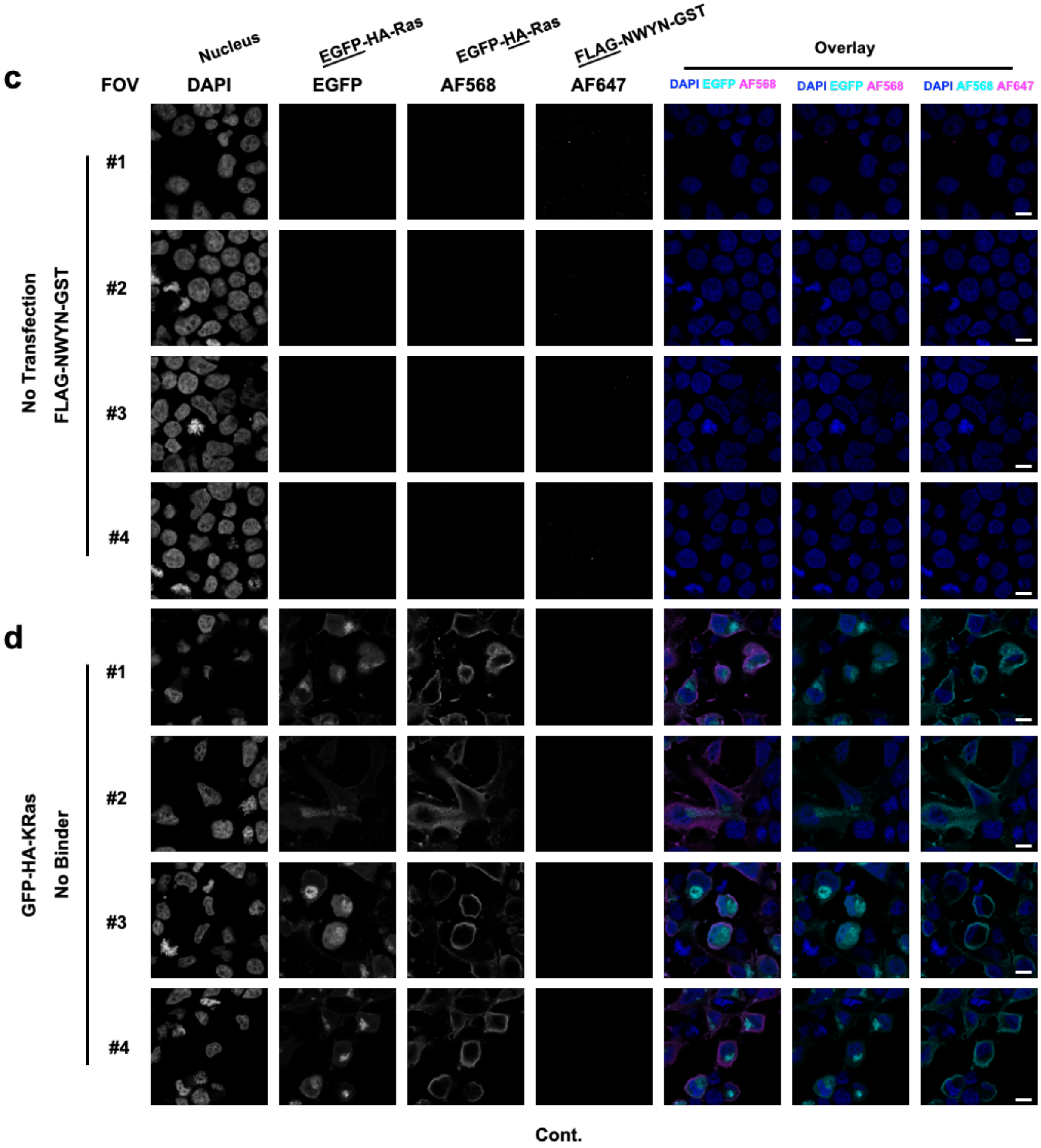


**Supplementary Fig. 26** *(Continued)*

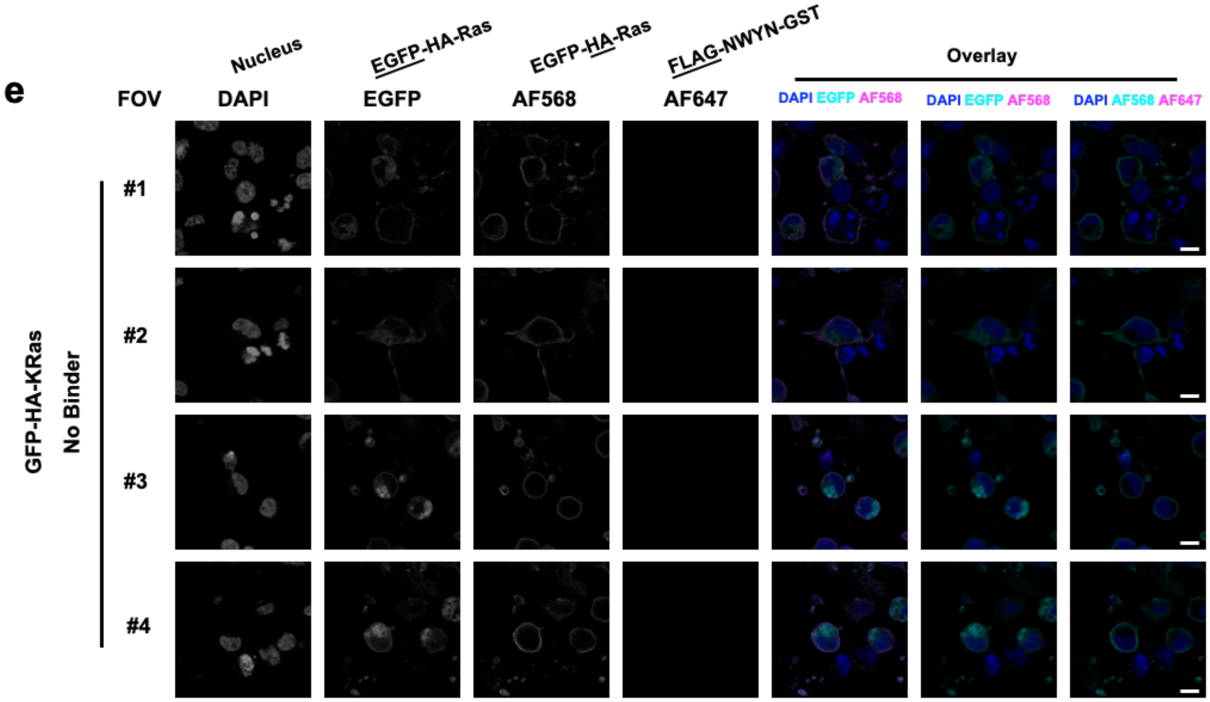


**Supplementary Fig. 26** *(Continued)*

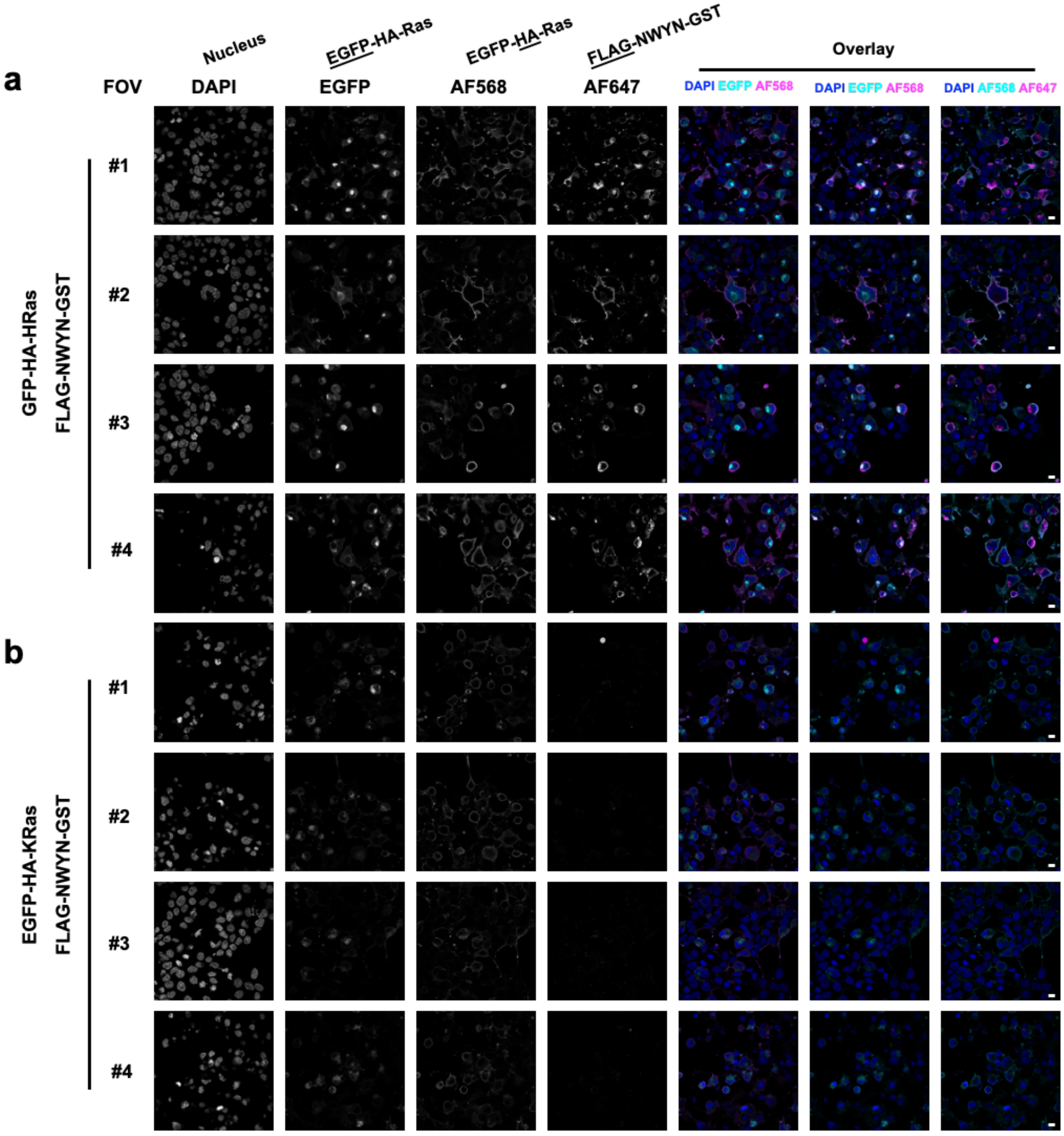


Cont.

**Supplementary Fig. 27: Zoom-out micrographs corresponding to 1× confocal zooms of the fields of view (FOVs) shown in Figure S26.** HEK293T cells overexpressing GFP-HA-HRas or GFP-HA-KRas fusion protein were stained with anti-HA/anti-rabbit-AF568, 3xFLAG-GST-NWYN/anti-FLAG/anti-Ms-AF647, and DAPI **(a,b)**. Untransfected and binder-omitted controls were included as background references. **(c-e)**. Four fields of views (FOVs) were shown per condition. *Top insets and underlined labels denote the intended viewing targets and corresponding color overlay schemes. Images were acquired using a 63× objective lens with 1× confocal zoom. Scale bar, 10 µm.* (*Continues on next page*)

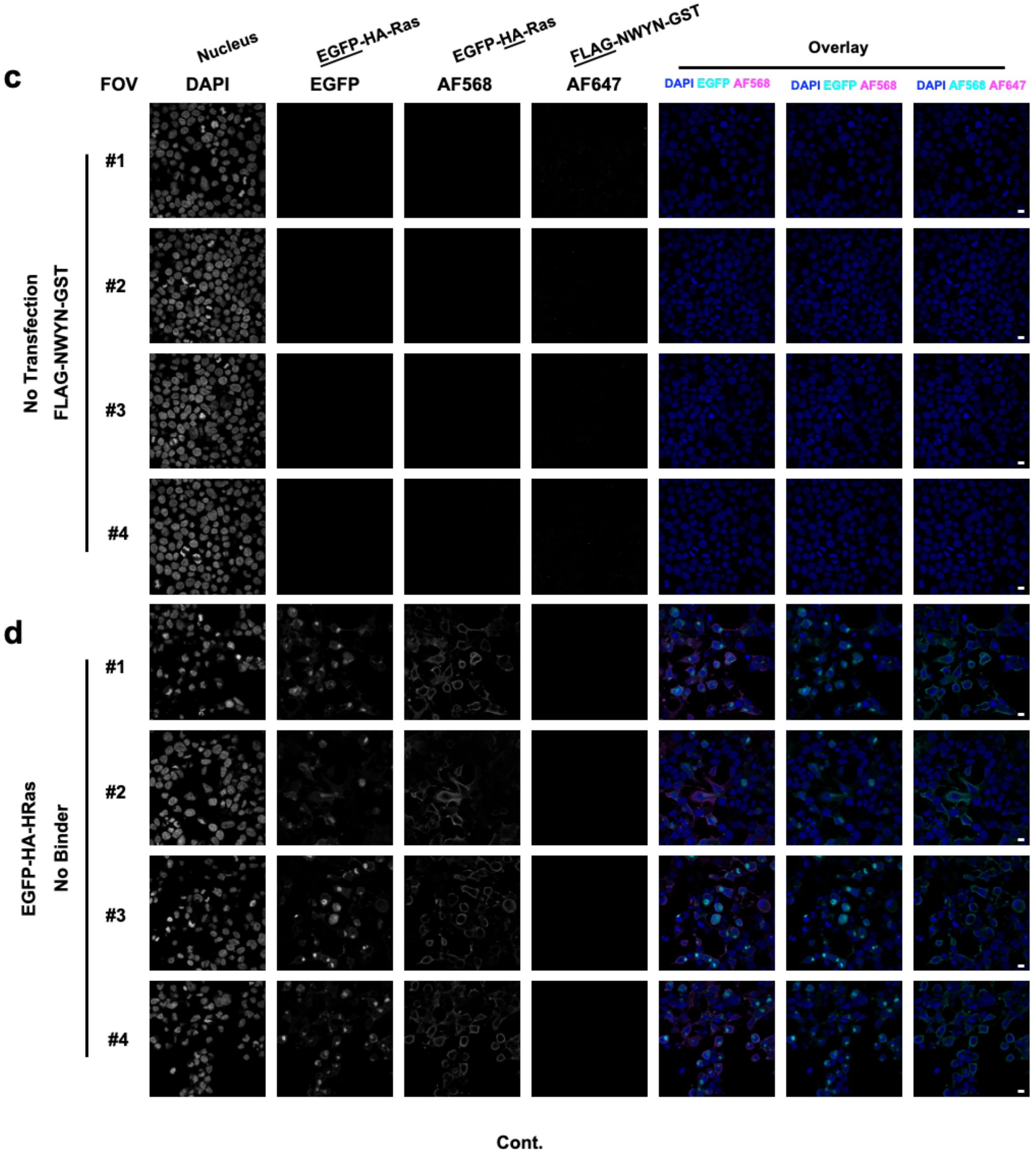


**Supplementary Fig. 27** *(Continued)*

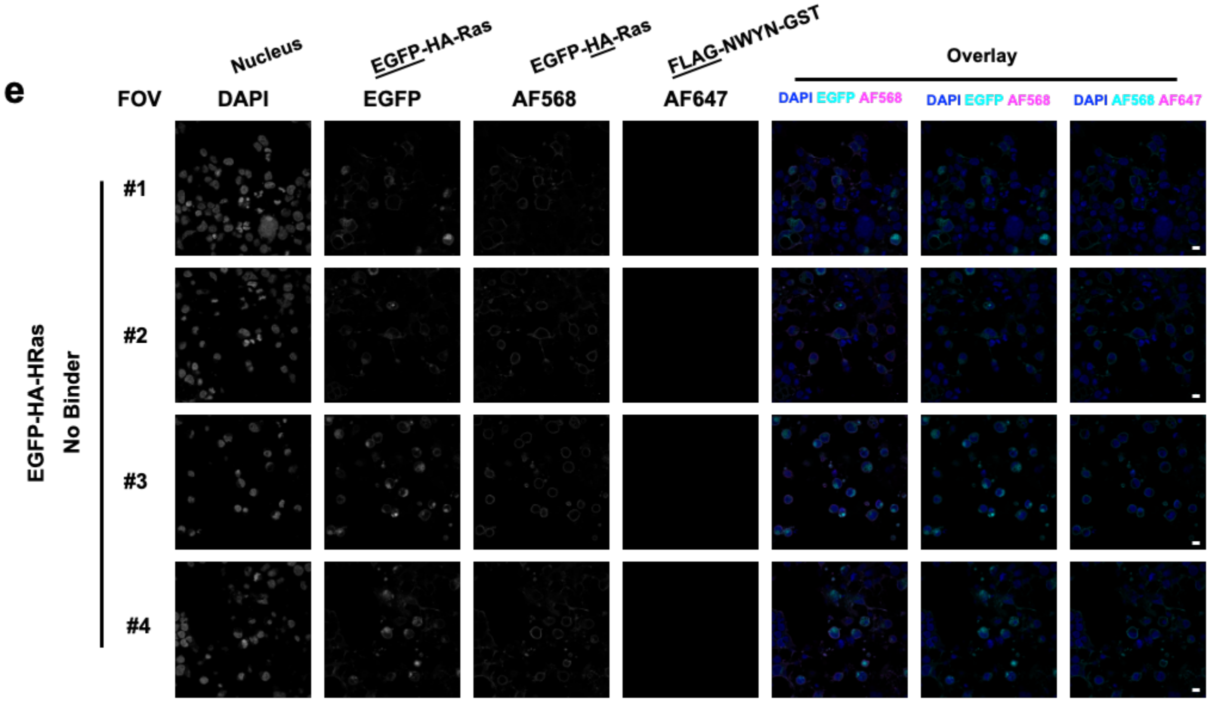


**Supplementary Fig. 27** *(Continued)*

### Additional Materials and Methods

#### Cloning and Bacterial Strain Handling

All plasmids and phage were cloned by Gibson Assembly (GA) of PCR fragments generated using Q5 DNA polymerase (NEB). All primers were ordered from IDT. For plasmids, GA mixtures were transformed into chemically competent DH10β *E. coli* and after a 1 h outgrowth in 2xYT media, were plated on antibiotic selective agar plates to isolate individual clones. For phage, GA mixtures were transformed into chemically competent S1030-1059 *E. coli,* and after a 2 h outgrowth, a plaque assay was performed to isolate individual phage clones. All plasmids and phage were confirmed by Sanger sequencing. All plasmid maps with annotations of key features are available in **Supplementary Table 6**. For constructing selection (+AP/-AP) and *E. coli* luciferase (2-22/N-lux/C-lux) strains, S1030 *E. coli* was made chemically competent and then single or double transformations were used (and then repeated as needed until all plasmids were incorporated). *E. coli* strains were grown on agar plates static at 37 °C or in solution at 37 °C with 200 rpm shaking with Luria Broth (LB) supplemented with the appropriate antibiotic unless otherwise indicated. Antibiotics were used at standard concentrations: kanamycin (40 ug/mL), chloramphenicol (33 ug/mL), and carbenicillin (100 ug/mL).

#### Plaque Assays

Activity independent plaque assays can be used to determine the phage titer via plaque counting. Activity dependent plaque assays can be used to check for robust phage replication on a given strain. For activity independent plaque assays, an S1030-1059 *E. coli* culture (1059 plasmid encodes *gIII* expressed from the phage shock promoter to produce *gIII* after phage infection), is grown to stationary phase in LB with carbenicillin, subcultured 1/10 in fresh LB with antibiotic to an OD600 of 0.4-0.6, and then used as the selection strain in the plaque assay. Similarly, for activity dependent strains, S1030 with a +AP (and -AP) were grown similarly for use in the plaque assay. For the plaque assay, an initial dilution of the stock can be added based on the expected titer, but generally, 2 µL of a phage stock (or diluted stock) is added to 100 µL of subculture, mixed, and then serially diluted (2 µL into 100 µL) to create 4 dilutions. 750 µL of 50 °C top agar (7 g/L agar, 25 g/L LB) was added to each dilution and then transferred in its entirety to one quadrant of a bottom agar plate (15 g/L agar, 15 g/L LB). After 10-16 h of incubation at 37 °C, plaques become visible and were counted in the quadrant with 10-200 plaque forming units (PFU).

#### Split-RNAP *E. coli* Luciferase Assays

We followed our previously reported[1] protocol for our split RNA polymerase *E. coli* luciferase assay (schematic shown in **Supplementary Fig. 1**). Luciferase strains were grown overnight in LB with appropriate antibiotics and then subcultured 7.5 µL into 143 µL of LB with kanamycin, chloramphenicol, carbenicillin, and L-arabinose (2 mg/mL final concentration) in white side, clear bottom 96-well assay plates (Corning 3610) and incubated at 37 °C with shaking (200 rpm) for 3.5 h prior to reading the OD600 and luminescence signal on a BioTek Synergy Neo2 plate reader. Luminescence signal is normalized by OD600 and all signals are compared as the fold-change to the non-binding affibody signal.

#### NGS of PANCS Hits

We used the Amplicon EZ service provided by Genewiz (Azenta) for Illumina sequencing of each of our hits (GENEWIZ from Azenta — Amplicon-EZ) which provides 50,000+ paired end reads per sample. We used primers (**Supplementary Table 7**) to install Illumina partial adaptors and barcodes to PCR products extending from the linker to after the stop codon of our scaffold in phage. Qubit dsDNA High Sensitivity kit is used to determine an accurate concentration of the sample prior to dilution and submission for sequencing. BB Merge was used to merge each paired end reads (https://jgi.doe.gov/data-and-tools/software-tools/bbtools/bb-tools-userguide/ bbmerge-guide/) and then MatLab was used to separate reads by barcode and to translate reads using modified scripts as described previously (MatLab scripts provided as a supplement).[1]

### New Library Construction

We utilize our previously reported $10^8$ affitin, $10^8$ affibody, and $10^{10}$ affibody libraries for this work.[1] We followed the general protocol for library construction that we previously reported[1] with the following modifications:

*Monobody Library*: The monobody library was cloned from Monobody (hSUMO1) phage (Supplementary Table 6), using MS-785 and MS-786 primers (Supplementary Table 7) by Q5 DNAP with GC enhancer with a $T_a$ of 71 °C. A ~$1*10^7$ library was generated following the general protocol with one 3 mg transformation.

*Nanobody Library*: The nanobody library was cloned from Nanobody (gfp) phage (Supplementary Table 6), using MS-802 and MS-803 primers (Supplementary Table 7) by Q5 DNAP with GC enhancer with a $T_a$ of 68 °C. A ~$5*10^7$ library was generated following the general protocol (except with PstI-HF was the restriction enzyme rather than NheI-HF) with one 3 mg and one 5 mg transformation, pooled together for the library outgrowth to make a single nanobody library.

### Protein Purification

Protein purification was completed as previously described[1] using C-terminally His-tagged Target proteins (HRAS (1-169), KRAS (1-166), and NRAS (1-166) and binder as 3xFLAG – GST- binder fusions.

### Surface Plasmon Resonance

Surface Plasmon Resonance was performed on a Biacore 8000 using a NTA chip for immobilizing the His-tagged target proteins. Target concentrations were optimized to elicit a response of ~50-100 RU (180 s of 5 uL/s) and then a range of binder concentrations were tested to identify concentrations that produced robust binding (90 s of 30 uL/s). All SPR conducted at 10 °C to maintain slow dissociation of the His-tagged immobilized protein. All dose-responses were fit to a kinetic model for 1:1 binding using the Biacore evaluation software – all fits passed the quality checks in this software (**Supplementary Figure 11** and **Supplementary Table 3**).

### Split Nano-Luciferase Assay

62.5 ng of the N-terminus of Nano-Luciferase fused to binder plasmid and 62.5 ng of the RAS fused to the C-terminus of Nano-Luciferase plasmid were co-transfected into HEK293T cells using Lipofectamine 2000 in clear 96 well plates (Corning 3595). Transfection was performed in triplicate. After 36 hours, the Nano-luciferase activity was measured using Nano-Glo® Live Cell Assay System (Promega, N2011).

**Supplementary Table 6: Plasmids**

**Phage**

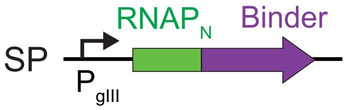


| ID | Description | Binder AA Sequence | Plasmid map |
|---|---|---|---|
| 70-70 | Gen 2 RNAP$_N$-RAF (WT) | SKTSNTIRVFLPNKQRTVVNVRNGMSLHDCLMKALKVRGLQPECCAVFRLLHEHKGKKARLDWNTDAASLIGEELQVDFL | https://benchling.com/s/seq-JJq1S2iNyzsJmd6xiEdv?m=slm-aRoQFIVy6IMYBSH2fwfJ |
| 70-81 | Gen 2 RNAP$_N$-Affibody (RAF) | VDNKFNKEVNLAADEIWLLPNLNNQQAWAFITSLKDDPSQSANLLAEAKKLNDAQAPK | https://benchling.com/s/seq-jlkztBe79lrkgW3fBhwO?m=slm-HWw47rRT6Gkt6WjA4nRo |
| 75-103 | Gen 2 RNAP$_N$-Affibody (PDL1) | VDNKFNKEPRAARLEITVLPNLNREQGGAFIVSLWDDPSQSANLLAEAKKLNDAQAPK | https://benchling.com/s/seq-w0kNAOKhSETKaNFpG3r2?m=slm-ZXJq3YteiKU5l3VNQVs7 |
| 70-78 | Gen 2 RNAP$_N$-Affitin (SasA) | GSVKVKFVVRGEEKEVDTSKIRSVWRGGKWVDFSYDDNGKQGYGYVLEKDAPKELLDMLARAEREKKL | https://benchling.com/s/seq-kGbU1x3dZtABiwGPTOvT?m=slm-ktQkaNLZJpvR0y70HFcJ |
| 70-80 | Gen 2 RNAP$_N$-Monobody (hSUMO1) | VSSVPTKLEVVAATPTSLLISWDAPAVTVDHYVITYGETGGNSPVQEFTVPGSKSTATISGLSPGVDYTITVYAWVSDEYTHGYYSYSPISINYRT | https://benchling.com/s/seq-BMAv02BGSfvUsKloWoMq?m=slm-k4Z8qtY23ikBzE9EvwAc |
| 70-76 | Gen 2 RNAP$_N$-Nanobody (GFP) | QVQLVESGGALVQPGGSLRLSAAASGFPVNRYSMRWYRQAPGKEREWVAGMSSAGDRSSYEDSVKGRFTISRDDARNTVYLQMNSLKPEDTAVYYVNVNVGFEYWGQGTQVTVSS | https://benchling.com/s/seq-lZcz30RbE2ALQNc1vqrv?m=slm-d6DGBULmHItVoc5Rur9f |

**+APs**

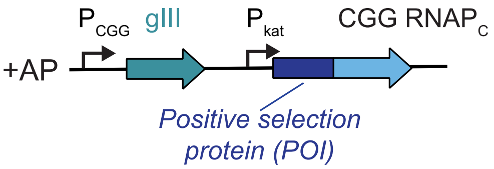


| ID | Target | ORI, RBS (P$_{CGG}$)/(P$_{kat}$) | Plasmid Map |
|---|---|---|---|
| 75-136 | KRAS | p15a, SD8/SD8; 60 AA linker | https://benchling.com/s/seq-ZeZ4aYcPMa4jt6kE0JA8?m=slm-urQbtWwArEqCnNkcetG1 |
| 75-124 | NRAS | p15a, SD8/SD8; 60 AA linker | https://benchling.com/s/seq-rBJD0ntLGgEuAuq1kSED?m=slm-yshqst61vCjixpRdgNbY |
| 75-135 | HRAS | p15a, SD8/SD8; 60 AA linker | https://benchling.com/s/seq-GOe4zjTl1Mdf4PRb31OS?m=slm-aVc4rYGqhQtzmXMU7JX5 |
| 75-126 | LC3B | p15a, SD8/SD8; 60 AA linker | https://benchling.com/s/seq-BVnxoGkEIF8WqzyciCqM?m=slm-zsHlH0xb2vn5ws0dScqE |

### -APs

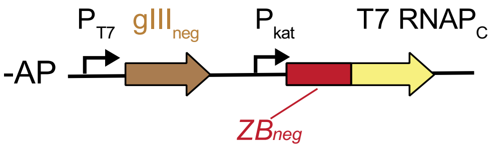


| ID | Description | Plasmid map |
|---|---|---|
| 73-79 | $ZB_{neg}$, 60 AA linker, sd8/sd8 | https://benchling.com/s/seq-qptnks4V9vCAkgQsGtmf?m=slm-0SHHTsCgDhaxvivYq6yY |
| 80-158 | KRAS wt, 60 AA linker, sd8/sd8 | https://benchling.com/s/seq-CnIHVxecMuV7hhFWqUuV?m=slm-UKodik5QvSD7wLE0jPdj |
| 73-97 | LC3B wt, 60 AA linker, sd8/sd8 | https://benchling.com/s/seq-KUpjN2HO7lldRM2KeqRK?m=slm-uVptyi8rGUJDCbtsXOul |
| 73-107 | LC3B LIR Mutant, 60 AA linker, sd8/sd8 | https://benchling.com/s/seq-gdPEdNrdmZ5Ly0GrvklI?m=slm-L20S0imALO86u7Oy3gvH |

### Plasmids for luciferase assay

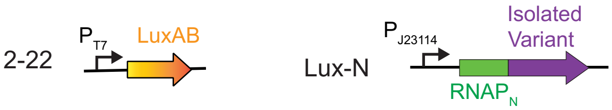


| ID | Description | AA Sequence | Plasmid map |
|---|---|---|---|
| 2-22 | $P_{T7}$ driven LuxAB on pSC101 | | https://benchling.com/s/seq-Udyz6dlESk9lOHBzqfsO?m=slm-pPVnaTxvld7HclZ8lnVO |
| 77-153 | Lux-N Affibody (PDL1) | VDNKFNKEPRAARLEITVLPNLNREQGGAFIVSLWDDPSQSANLLAEAKKLNDAQAPK | https://benchling.com/s/seq-4Hat7A1iLPOjVB5SfuyL?m=slm-kG73luQP4lmO9k1o5Xk0 |
| 77-154 | Lux-N Affibody (RAF) | VDNKFNKEVNLAADEIWLLPNLNNQQAWAFITSLKDDPSQSANLLAEAKKLNDAQAPK | https://benchling.com/s/seq-BbfsAlyrXYrH5iGF0TXP?m=slm-8PKvi1so0qMW70ijQh4W |
| General Subcloned Library Variant Library Variant | | Affibody Example VDNKF?KE???A???I??LPNLN??Q??AF??SL??DPSQSANLLAEAKKLNDAQAPK | https://benchling.com/s/seq-mVSKERuVdRGIeQ7j4OKo?m=slm-Hk4Rjrf5ZYCffF4uTrbZ |

### Lux-C

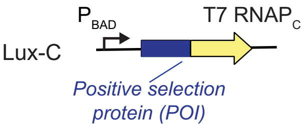


| Lux-C | Target | Lux-C Map |
|---|---|---|
| 76-97 | NRAS | https://benchling.com/s/seq-MNEcNbqEA5bJjd5OxWp4?m=slm-kItXGR5rmeQg46qZz5G8 |
| 76-98 | HRAS | https://benchling.com/s/seq-pgTuUgg65DUpR1fn20uo?m=slm-zeyRxnr9T8TrgjagcSCj |
| 76-96 | KRAS | https://benchling.com/s/seq-YNF06s3yMNWhoWHllQZm?m=slm-pOpYJTkPYLfYd25kMJqP |
| 76-77 | GABARAP | https://benchling.com/s/seq-QWrUgcltA0dgKyKnI5fN?m=slm-TRc3OoEsRBgRJn0jMb2z |
| 76-79 | LC3B | https://benchling.com/s/seq-rlt5IzSBDshWhl9Rb2fu?m=slm-SμL56BpmJVP1Nz5nKQcW |
| Any mutant versions of these plasmids are mutants from the parent plasmid listed here. | | |

### pET for protein purification

| ID | Description | Plasmid map |
|---|---|---|
| 79-123 | KRAS WT 1-169 pET28a (HIS tag) | https://benchling.com/s/seq-YblODMQQm4pwkHpmK8FP?m=slm-EtvwnaIhlQmk8L0MUFM4 |
| 79-16 | HRAS 1-166 pET28a (HIS tag) | https://benchling.com/s/seq-S9QGfmsHabG5a2TEJS5d?m=slm-WKYQHvChRXtMwKTw4Pj8 |
| 79-17 | NRAS 1-166 pET28a (HIS tag) | https://benchling.com/s/seq-vNzxovEXiWWG7wpnU9pQ?m=slm-yavXBhOeFZgrTpL2o1K8 |
| 80-171 | Example Affibody Library Binder pET30 (GST tag) | https://benchling.com/s/seq-Uklg7K2IUQhI4TbmX6P3?m=slm-YCHStj8V8OZv5zNbd8rf |

**CMV Plasmids**

| ID | Description | Plasmid Map |
|---|---|---|
| 80-101 | Target split NanoLuc | https://benchling.com/s/seq-JCCd6CqbklNPQebimKmJ?m=slm-XUDQtXptgrmaXxwQvGvt |
| | Binder split NanoLuc | https://benchling.com/s/seq-kHb13xPqEKHeQ332Pchh?m=slm-tNeDLDnlXlavuDRaNGkD |
| 81-16 | Affibody Expression Plasmid (HA tag) | https://benchling.com/s/seq-AiR0IfgsQPlabGw7lv7q?m=slm-yc3nhQDr9K6FVAXvg3k9 |

**Supplementary Table 2: Primers**

| ID | Description | **Sequence** ($X_{12}$ indicates sequence specific to target) |
|---|---|---|
| VC-525 | Forward primer for qPCR | ggttcagcaaggtgatgctt |
| VC-526 | Reverse primer for qPCR | accgcctcacctctgtttta |
| MS-785 | Forward primer for Monobody Library | CATCATCATgctagcCCNGGNGTNGACTACACNATH<br>ACNGTNTATGCNTGGRSYDVYNNBRRYNNBNNBN<br>NBNNBNNBKMYAATTATTCGCCGATTTCTATCAAC<br>TATCGTACCTAATGGAGATTTTCAACATGCTCCCT<br>CAATCGG |
| MS-786 | Reverse primer for Monobody Library | CATCATCATgctagcGCCGGAAATAGTAGCRKWRK<br>WRKWRKWNCCCGGNACAGTAAATTCTTGNACCG<br>GNGAATTNCCNCCNGTTTCNCCGTANGTDATNAC<br>GTARYMVNNNGHVNNVNNVNNRKRRKHATCCCAA<br>GAAATCAGCAAGCTGGTCGGAGTTGCGGC |
| MS-802 | Forward primer for Nanobody Library | CATCATCATctgcagATGAATTCNTTAAAACCAGAAG<br>ATACNGCNGTNTATTATRYHNWYGTAHHYGTAGG<br>TNNYHHYTACNNBGGTCAAGGTACGCAGGTTACT<br>GTGTCCTCCTAATGGAG |
| MS-803 | Reverse primer for Nanobody Library | CATCATctgcagATANACNGTATTTTTNGCATTATCT<br>CTNGAAATNGTAAATCTTCCTTTNACNGAATCNGC<br>ATARWNAGTRDDRNNACCRNNAGARNNAATNSCN<br>GCNACCCATTCTCTTTCTTTNCCTGGNGCTTGAC<br>GATACCARWNCATRDDRNNRNNRNNCACCGGGA<br>AACCGCTCGCAGCC |
| | General Forward for insert for cloning scaffold into phage or Lux-N | tggctctggctctggctcgagcXXXXXXXXXXXX |
| | General Reverse for insert for cloning scaffold into phage | cgattgagggagcatgttgaaaatctccattaXXXXXXXXXXXX |
| | General Reverse for insert for cloning scaffold into Lux-N | gctgaggagtgccgttaattaagtttaXXXXXXXXXXXX |
| BR-305 | Forward for vector for cloning scaffold into phage | tggagattttcaacatgctccctcaatcg |
| MS-40 | Forward for vector for cloning scaffold into Lux-N | taaacttaattaacggcactcctcagcaaatataatgacc |
| KJ-14 | Reverse for vector for cloning scaffold into phage or Lux-N | CTCTGGCTCTGGCTCGAGC |
| | General Forward for insert for cloning target into +AP | ggatccctcgaaaggaggaaaaaaaaATGXXXXXXXXXXXX |
| | General Forward for insert for cloning target into Lux-C | gtctgacataaatgaccgctATGXXXXXXXXXXXX |
| | General Reverse for insert for cloning target into +AP/Lux-C | ctttaccgcttccacccgacgtXXXXXXXXXXXX |
| KJ-19 | Forward for vector for cloning target into +AP or Lux-C | ACGTCGGGTGGAAGCGGT |
| MS-46 | Reverse for vector for cloning target into +AP | tttttttcctcctttcgagGGATCCtaggtag |
| BR-116 | Reverse for vector for cloning scaffold into Lux-C | ACAACTCAAGTCTGACATAAATGACCGCT |
| MS-659 | Forward for insert for subcloning library variants into Lux-N | cacgtctacaagGGTGGCTCTG |
| JD-856 | Reverse for insert for subcloning library variants into Lux-N | tcaatcggttgaatgtcgccctt |

| | | |
|---|---|---|
| MS-660 | Forward for vector for subcloning library variants into Lux-N | aatcggttgaatgtcgcccttacttaattaacggcactcctcagc |
| MS-661 | Reverse for vector for subcloning library variants into Lux-N | cacgtctacaagGGTGGCTCT |
| MS-1156 | Reverse primer for Amplicon-EZ NGS for all barcodes (purple is adaptor, green is priming region) | gactggagttcagacgtgtgctcttccgatctgggcgacattcaaccgatt gaggg |
| MS-1144 | Forward primer for Amplicon-EZ NGS Barcode 1 (red is adaptor, blue is barcode, green is priming region) | acactctttccctacacgacgctcttccgatctAAGGTTgcgcgtagggc acgtctacaag |
| MS-1145 | Forward primer for Amplicon-EZ NGS Barcode 2 | acactctttccctacacgacgctcttccgatctTAGGATgcgcgtagggc acgtctacaag |
| MS-1146 | Forward primer for Amplicon-EZ NGS Barcode 3 | acactctttccctacacgacgctcttccgatctTGAGATgcgcgtagggc acgtctacaag |
| MS-1147 | Forward primer for Amplicon-EZ NGS Barcode 4 | acactctttccctacacgacgctcttccgatctAGTGATgcgcgtagggc acgtctacaag |
| MS-1148 | Forward primer for Amplicon-EZ NGS Barcode 5 | acactctttccctacacgacgctcttccgatctAAGTGTgcgcgtagggc acgtctacaag |
| MS-1149 | Forward primer for Amplicon-EZ NGS Barcode 6 | acactctttccctacacgacgctcttccgatctAAGTTGgcgcgtagggc acgtctacaag |
| MS-1150 | Forward primer for Amplicon-EZ NGS Barcode 7 | acactctttccctacacgacgctcttccgatctAGATGTgcgcgtagggc acgtctacaag |
| MS-1151 | Forward primer for Amplicon-EZ NGS Barcode 8 | acactctttccctacacgacgctcttccgatctAGTAGTgcgcgtagggc acgtctacaag |
| MS-1152 | Forward primer for Amplicon-EZ NGS Barcode 9 | acactctttccctacacgacgctcttccgatctGAATTGgcgcgtagggc acgtctacaag |
| MS-1153 | Forward primer for Amplicon-EZ NGS Barcode 10 | acactctttccctacacgacgctcttccgatctGATATGgcgcgtagggc acgtctacaag |
| MS-1154 | Forward primer for Amplicon-EZ NGS Barcode 11 | acactctttccctacacgacgctcttccgatctGTAATGgcgcgtagggc acgtctacaag |
| MS-1155 | Forward primer for Amplicon-EZ NGS Barcode 12 | acactctttccctacacgacgctcttccgatctGTATAGgcgcgtagggc acgtctacaag |
| MS-1346 | Forward primer for Amplicon-EZ NGS Barcode 13 | acactctttccctacacgacgctcttccgatctAACCTTgcgcgtagggc acgtctacaag |
| MS-1347 | Forward primer for Amplicon-EZ NGS Barcode 14 | acactctttccctacacgacgctcttccgatctTACCATgcgcgtagggc acgtctacaag |
| MS-1348 | Forward primer for Amplicon-EZ NGS Barcode 15 | acactctttccctacacgacgctcttccgatctTCACATgcgcgtagggc acgtctacaag |
| MS-1349 | Forward primer for Amplicon-EZ NGS Barcode 16 | acactctttccctacacgacgctcttccgatctACTCATgcgcgtagggc acgtctacaag |
| MS-1350 | Forward primer for Amplicon-EZ NGS Barcode 17 | acactctttccctacacgacgctcttccgatctAACTCTgcgcgtagggc acgtctacaag |
| MS-1351 | Forward primer for Amplicon-EZ NGS Barcode 18 | acactctttccctacacgacgctcttccgatctAACTTCgcgcgtagggc acgtctacaag |
| MS-1352 | Forward primer for Amplicon-EZ NGS Barcode 19 | acactctttccctacacgacgctcttccgatctACATCTgcgcgtagggc acgtctacaag |

| | | |
|---|---|---|
| MS-1353 | Forward primer for Amplicon-EZ NGS Barcode 20 | acactctttccctacacgacgctcttccgatctACTACTgcgcgtagggcacgtctacaag |
| MS-1354 | Forward primer for Amplicon-EZ NGS Barcode 21 | acactctttccctacacgacgctcttccgatctCAATTCgcgcgtagggcacgtctacaag |
| MS-1355 | Forward primer for Amplicon-EZ NGS Barcode 22 | acactctttccctacacgacgctcttccgatctCATATCgcgcgtagggcacgtctacaag |
| MS-1356 | Forward primer for Amplicon-EZ NGS Barcode 23 | acactctttccctacacgacgctcttccgatctCTAATCgcgcgtagggcacgtctacaag |
| MS-1357 | Forward primer for Amplicon-EZ NGS Barcode 24 | acactctttccctacacgacgctcttccgatctCTATACgcgcgtagggcacgtctacaag |
